# Leaf form diversification in an heirloom tomato results from alterations in two different *HOMEOBOX* genes

**DOI:** 10.1101/2020.09.08.287011

**Authors:** Hokuto Nakayama, Steven D. Rowland, Zizhang Cheng, Kristina Zumstein, Julie Kang, Yohei Kondo, Neelima R. Sinha

## Abstract

Domesticated plants and animals display tremendous diversity in various phenotypic traits and often this diversity is seen within the same species. Tomato (*Solanum lycopersicum*; Solanaceae) cultivars show wide variation in leaf morphology, but the influence of breeding efforts in sculpting this diversity is not known. Here, we demonstrate that a single nucleotide deletion in the homeobox motif of *BIPINNATA*, which is a *BEL-LIKE HOMEODOMAIN* gene, led to a highly complex leaf phenotype in an heirloom tomato, Silvery Fir Tree (SiFT). Additionally, a comparative gene network analysis revealed that reduced expression of the ortholog of *WUSCHEL RELATED HOMEOBOX 1* is also important for the narrow leaflet phenotype seen in SiFT. Phylogenetic and comparative genome analysis using whole-genome sequencing data suggests that the *bip* mutation in SiFT is likely a *de novo* mutation, instead of standing genetic variation. These results provide new insights into natural variation in phenotypic traits introduced into crops during improvement processes after domestication.

## Main

Domestication and subsequent improvement processes have made animals and plants more suitable for agriculture and achieved improvement in their usability, quality, and yield (1). In contrast to domestication, usually occurring once for many crops, selection for improvement happened multiple times and in numerous locations, leading to varieties adapted to local conditions and needs (2). Consequently, many crops show fascinating morphological diversity. Indeed, Darwin focused on this morphological diversity more than 150 years ago and postulated that knowledge of the mechanisms underlying diversity generated under human selection would provide general principles for understanding the process of evolution under natural selection (3). Domesticated tomato, *Solanum lycopersicum* L. (Solanaceae), is one of the most economically important vegetable crops in the world (4). The domesticated tomato exhibits tremendous morphological variation because of a long breeding history (5).

Additionally, many heirloom tomatoes, varieties passed down through several generations within a family or specific regions, have an interesting breeding history predating 1940 when commercial hybrids first started becoming available. These heirloom cultivars, when compared to commercial tomatoes, show morphological variation and flavor profiles that are often favored by gardeners (6). Many heirloom tomatoes contain genetic loci that affect most of the target flavor chemicals to improve the flavor of modern commercial tomato (7), making heirloom tomatoes an interesting resource for research on tomato improvement. Silvery Fir Tree (SiFT) is a traditional Russian heirloom tomato (6). SiFT has a highly complex leaf phenotype, with leaflets that are narrower than those seen in processing tomatoes such as M82 (Figure 1A to 1D). Interestingly, SiFT is sometimes used as an ornamental and landscaping plant rather than a crop due to the unique leaf shape in this variety, although SiFT does produce edible fruit (6). However, the genetic basis underlying the unique leaf morphology and breeding history of this cultivar is still unknown.

**Figure 1.**
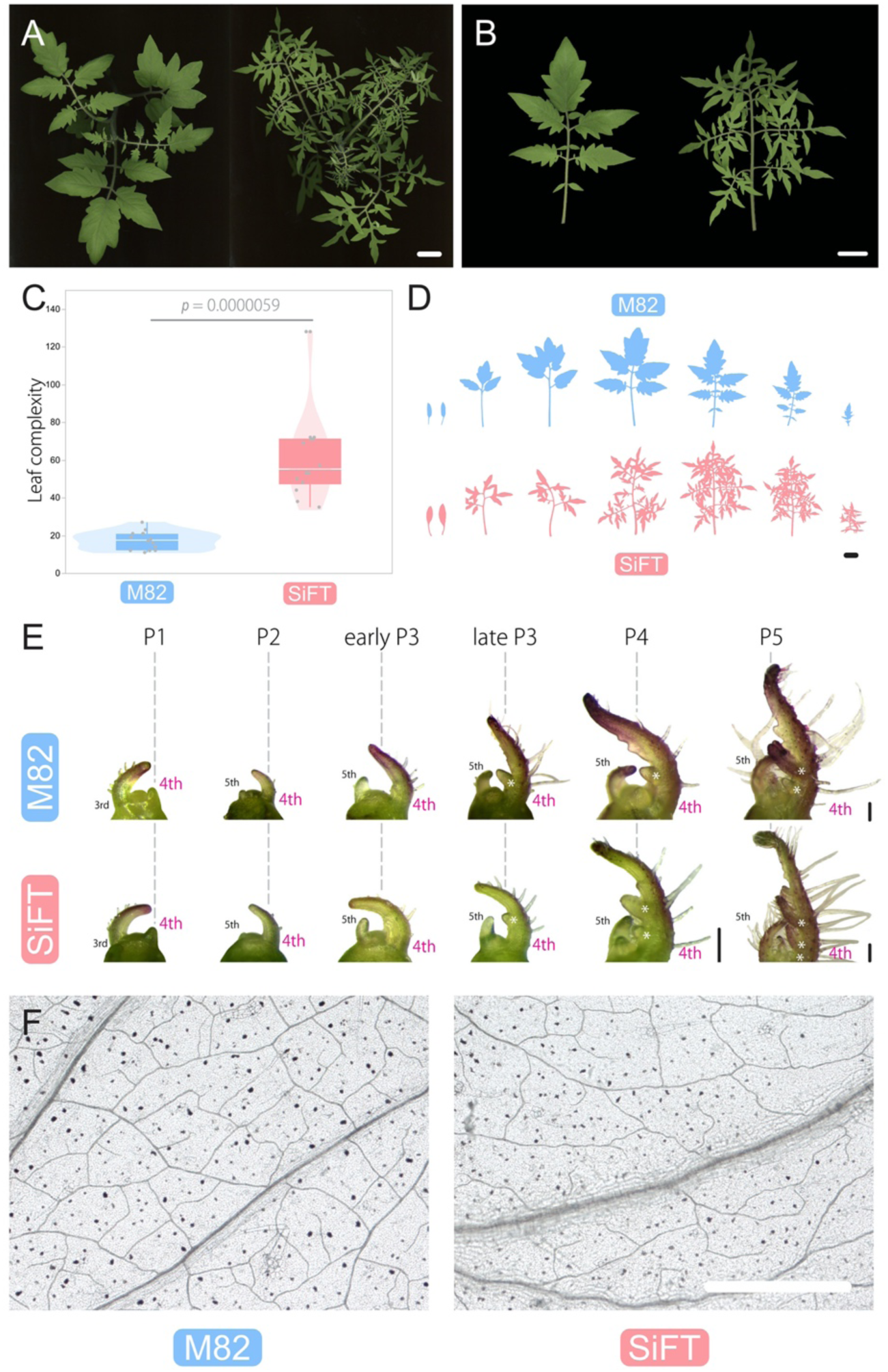
Gross morphology and development in M82 and SiFT leaves. Top view of shoots (A), and mature leaf morphology (B). The 4th leaves were used for (B). Left: M82; right: SiFT. (C) Comparison of leaf complexity (*N* = 14). *p* = 0.0000059 (Welch’s t-test). (D) Comparison of leaf morphology of M82 (upper) and SiFT (lower). All silhouettes are based on photographic images. The youngest leaf is at the right and the oldest (cotyledons) is at the left. (E) Developmental trajectory of M82 and SiFT leaf primordia. The 4th and 5th leaves were represented. (F) Cleared terminal leaflet images of M82 and SiFT. Bars = 2 cm in (A), (B) and (D), 100 μm in (E), and 1 mm in (F).

Here, we used a cross between SiFT and M82 to generate a mapping population and identified a single nucleotide deletion in the homeobox motif of a *BEL-LIKE HOMEODOMAIN* (*BELL*) gene, leading to a premature stop codon, and the highly complex leaf phenotype in SiFT. Based on genome sequencing, the *bip* mutation in SiFT is a *de novo* mutation that was not introgressed from other cultivars or wild species. Further, we use a combination of gene co-expression network analysis and CRISPR-Cas9 knockout mutants to show that reduced expression of the *WUSCHEL RELATED HOMEOBOX 1* (*WOX1*) ortholog is also important for the narrower leaflet phenotype and reduced leaf vascular density in SiFT. Additionally, we show that the classic tomato leaf mutation, *solanifolia*, is caused by mutations in the same *WOX1* gene. These results provide insights into natural variation in phenotypic traits introduced into heirloom tomatoes during improvement after domestication.

## Results

### SiFT has increased leaf complexity and reduced vascular density compared to M82

Leaf complexity (LC) in SiFT is higher than that in M82 (Fig. 1A to 1C). The observation of gross leaf morphology showed distinct morphological differences between M82 and SiFT starting from the 1st formed leaves (Fig. 1D). While leaf primordia from Plastochron1 (P1) to P3 stages are not strikingly different between the cultivars, from P4 stage onward, difference in the number of leaflet primordia are consistently observed between M82 and SiFT (Fig. 1E). Thus, SiFT leaf primordia at P4 and older stages are more active in generating leaflets compared to M82 leaves at the same developmental stage, and have a prolonged morphogenetic window compared to that in M82. Previous studies have shown leaf vascular density (LVD) variation among cultivars and mutants (8). Although no difference was observed in leaf anatomy around the midvein (Supplementary Fig. 1), LVD was different between M82 and SiFT (Fig. 1F). Therefore, SiFT differs from M82 in leaf complexity, developmental trajectory, and vascular density.

### SiFT has a mutation in a *BEL-LIKE HOMEODOMAIN* gene, *BIPINNATA*

To identify genes involved in the regulation of leaf complexity (LC), bulked segregant analysis (BSA) on an F2 population (198 individuals) derived from a cross between M82 and SiFT was utilized (Supplementary Fig. 2). Two phenotypically defined bulks showing difference in LC (High-LC bulk and Low-LC bulk; Supplementary Fig. 3) were used to detect a locus between 45000000 and 55000000 bp on chromosome 2 that controlled LC (Fig. 2A; top and Supplementary Fig. 4). Whole genome sequencing of SiFT was used to define sequence variants in the genome including the region defined by BSA. However, we detected more than 100 variants in this region in the SiFT genome (Fig. 2A; middle). To narrow down the number of candidates, we used Protein Variation Effect Analyzer (PROVEAN), which allows us to predict whether an amino acid substitution or indel has an impact on the biological function of a protein (Supplementary Fig. 2) (9). The PROVEAN analysis found only one deleterious variant in the region (Fig. 2A; bottom), located in the *BIPNNATA* (*BIP*: Solyc02g089940) gene (Fig. 2B), which is known to encode a BEL-LIKE HOMEODOMAIN (BLH) protein (10). *BIP* is located in the *BLH* clade in the *BELL/KNOX1* phylogeny (Supplementary Fig. 5). The 1 bp deletion at position 1674 within the homeobox domain generates a premature stop codon, and as a result the BIP protein is truncated in SiFT (Fig. 2B and 2C, and Supplementary Fig. 6). Additionally, we confirmed that the genome of a different SiFT accession, previously sequenced by Tieman and coworkers, also has the same *bip* mutation (7) (Supplementary Fig. 7).

**Figure 2.**
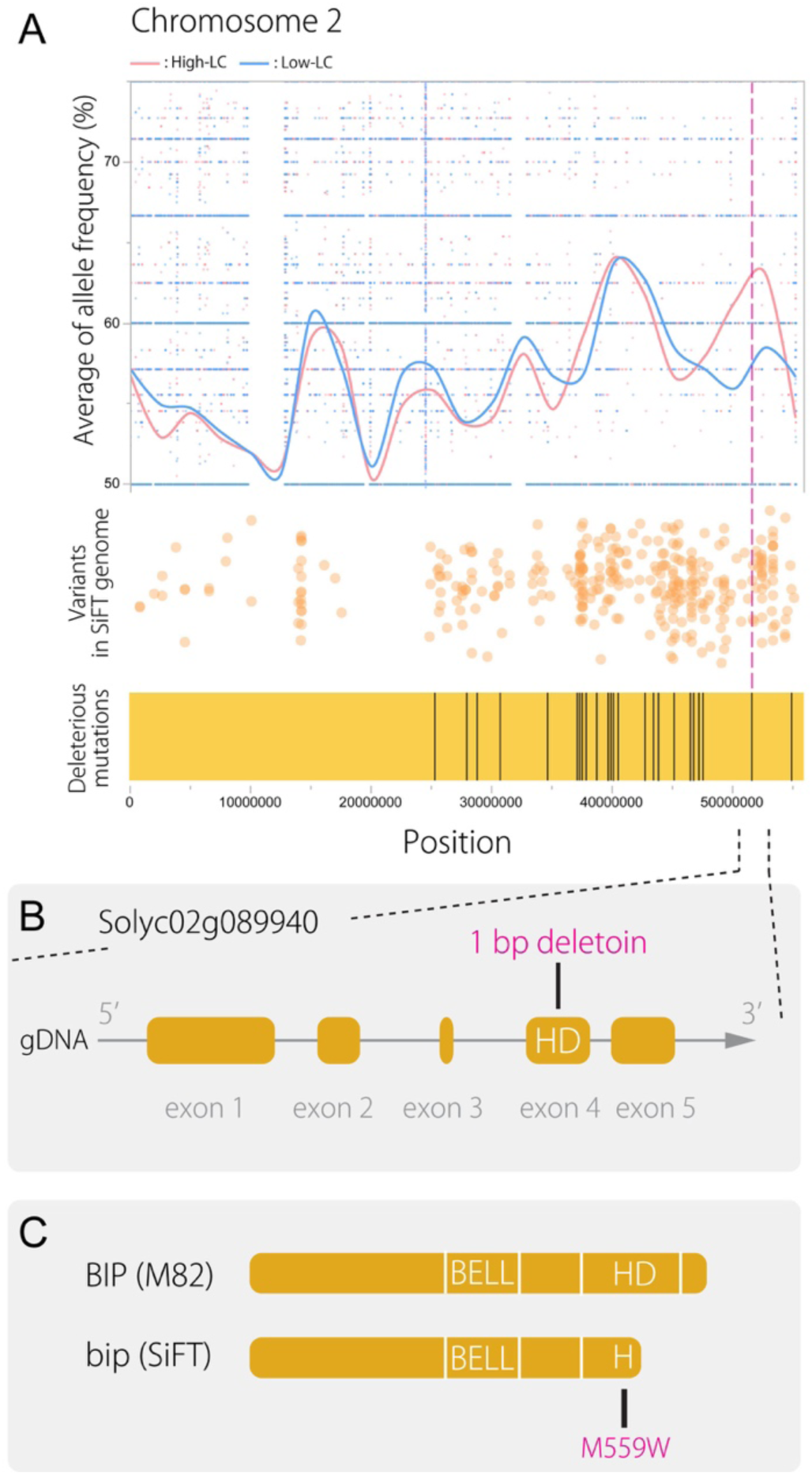
Identification of the causative mutation for the *BIPINNATA* gene. (A) Top: allele frequency between different pools of segregating populations (red: high complexity pool; blue: low complexity pool) is shown for chromosome 2 (Chr 2). Middle: variants (SNPs and indels) in SiFT from whole genome sequencing data. Each dot indicates variant position on Chr 2. Bottom: deleterious mutations in SiFT indicated from PROVEAN. Each vertical line indicates deleterious mutation on Chr 2. All panels (top, middle, and bottom) show the same scale on Chr 2. (B) Exon and intron structure of *BIPINNATA* (*BIP*). *BIP* gene contains five exons. SiFT contains an 1 bp deletion, which leads truncated protein and an amino acid change in the highly conserved amino acid of homeodomain (C).

**Figure 3.**
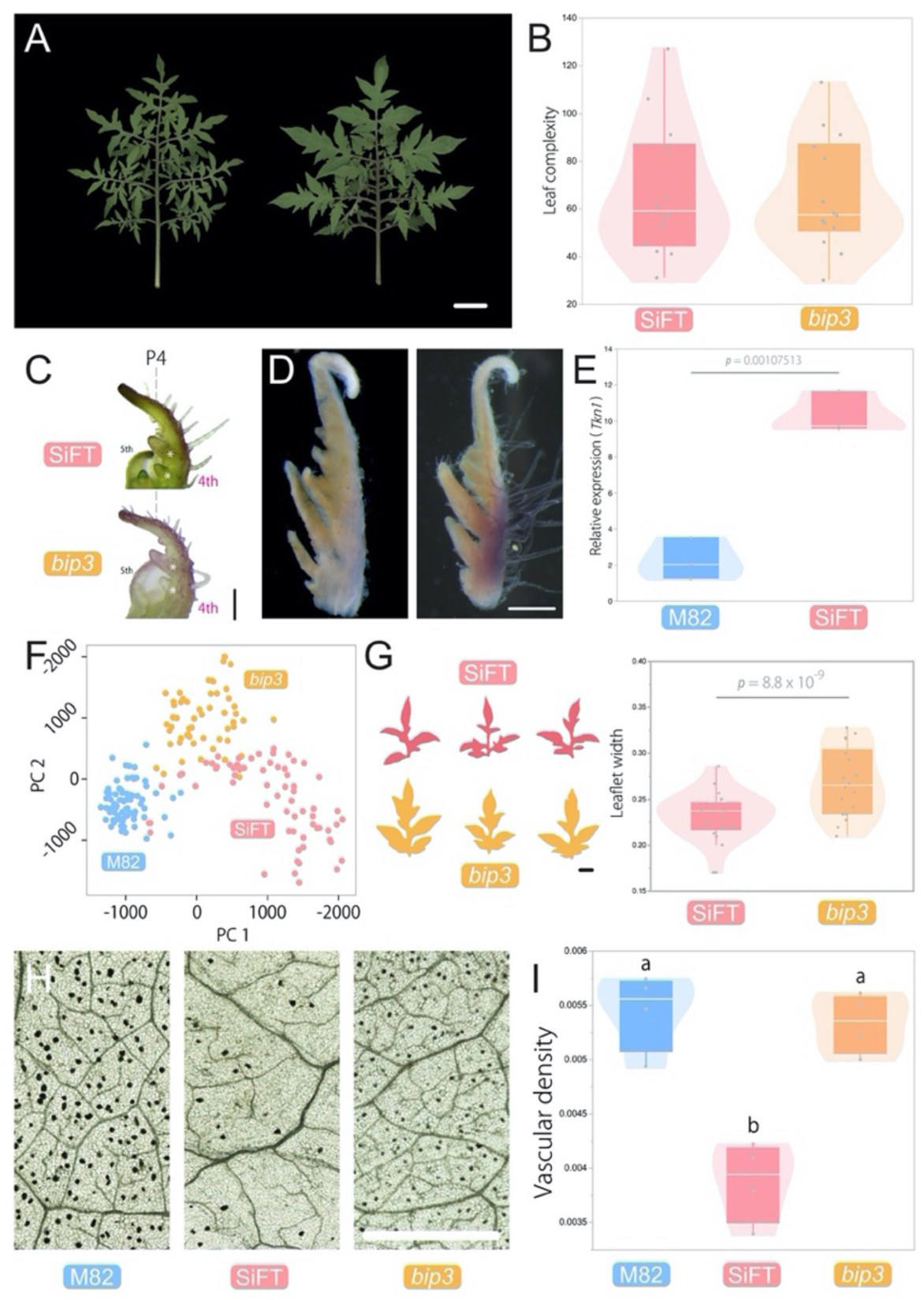
*bipinnata* leaf phenotypes. (A) Mature leaf morphology. The 4th leaves were used. Left: SiFT; right: *bip3*. (B) Comparison of leaf complexity (*N* = 14). (C) Leaf development of *bip3* at P4 stage. (D) Whole mount *in situ* localization of *BIP* transcripts in M82. Left: sense probe; Right: antisense probe. (E) Expression level of *Tkn1* in leaf primordia (*N* = 3). *p* = 0.00107513 (Welch’s *t*-test). (F) Deep learning-based nonlinear PCA with leaflet shapes (*N* < 55). Blue: M82, pink: SiFT, and orange: *bip3*. (G) Comparison of terminal leaflet morphology. Left; leaflet morphology used for leaf shape analysis. All silhouettes are based on scanned images. Right; results of leaf width measurement with terminal leaflets. *p* = 8.8 x 10^−8^ (Welch’s *t*-test). (H) Cleared terminal leaflet images of M82, SiFT and *bip3*. (I) Vascular density per unit area. The data was assessed using pair-wise comparisons with Tukey-Kramer HSD test. Bars = 2 cm in (A), 100 μm in (C), 500 μm in (D), 1 cm in (G), and 1 mm in (H).

**Figure 4.**
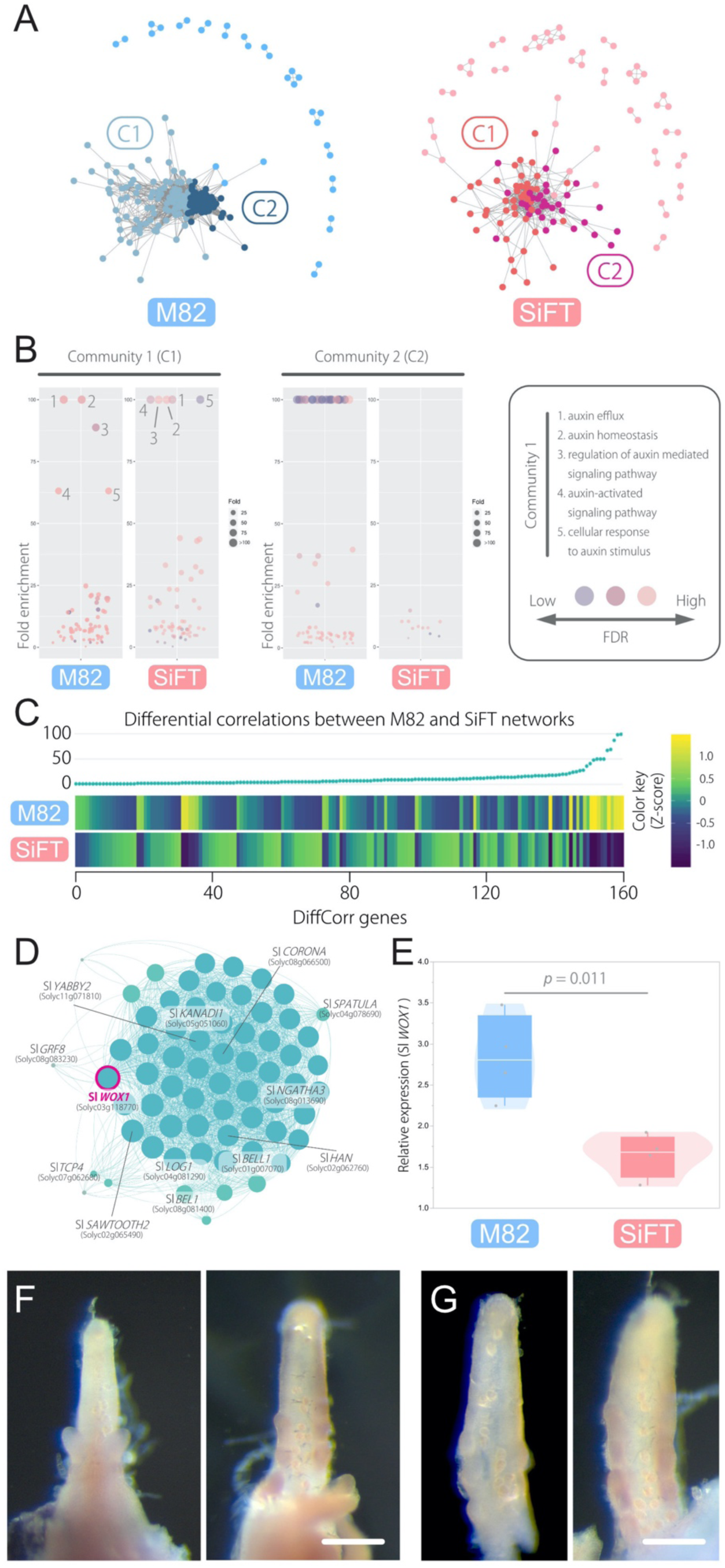
Gene co-expression network analysis with M82 and SiFT RNA-seq data. (A) Gene co-expression networks for genes involved in leaf development. Each node represents genes. Only nodes with at least one edge are represented. Left: M82; right: SiFT. (B) An overview of the enriched GO terms visualized by bubble plot. The analysis was performed by the community in each network (C1 and C2). Each bubble represents a GO term and only GO terms with higher Fold enrichment (>50) are represented. For full result of the GO enrichment analysis, please see Supplementary Table 3. (C) A profile of 160 DiffCorr genes. The plot on the top: the number of differential correlations of each DiffCorr gene. A higher number means more difference between M82 and SiFT networks. The heat map on the bottom: a comparison of expression level of each DiffCorr gene between M82 and SiFT. Each expression level is shown as a blue-to-yellow-colored scale. The 160 DiffCorr genes were sorted by the number of differential correlations (Left: low; right: high). The position of each gene is the same between the top and bottom panels. (D) The Sl *WOX1* gene network from M82 shown in (A). This network is consisted of genes only showing a direct connection to the Sl *WOX1*. (E) Expression level of Sl *WOX1* in leaf primordia (*N* = 4). *p* = 0.011 (Welch’s *t*-test). (F and G) Whole mount *in situ* localization of Sl *WOX1* transcripts in M82. (F) Leaf primordia. (G) Leaflet primordia. Left: sense probe; right: antisense probe in each panel. Bars = 100 μm in (F) and (G).

**Figure 5.**
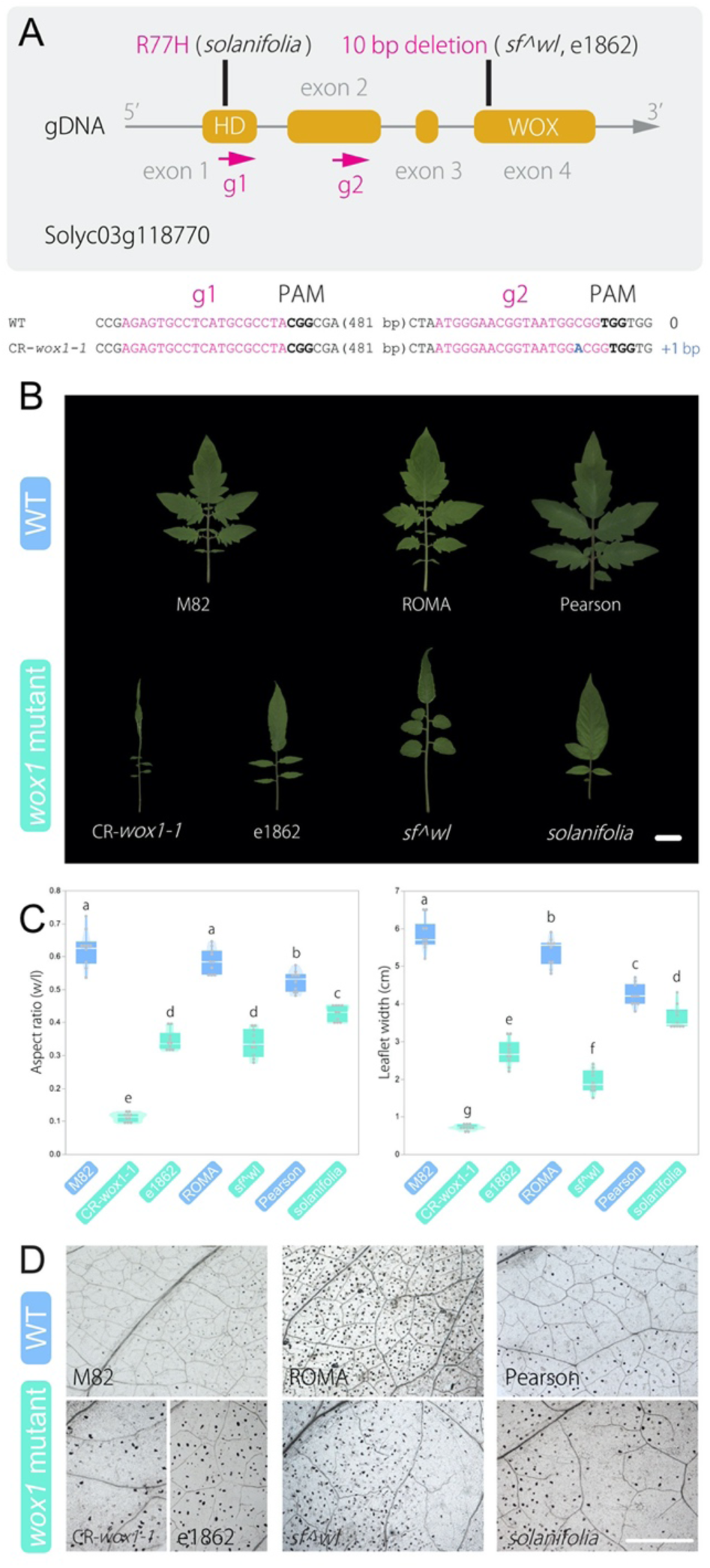
*solanifolia/slwox1* leaf phenotypes. (A) Exon and intron structure of *SF/SlWOX1*. The tomato *SF/SlWOX1* gene contains four exons. (B) Mature leaf morphology of *sf/slwox1* mutants. The 4th leaves were used. (C) Comparison of aspect ratio (width/length) and width of terminal leaflet (*N* = 10). Letters indicate significance groups; samples with the same letters are not significantly different. All data were assessed using pair-wise comparisons with Tukey-Kramer HSD test. (D) Comparison of vascular density. Cleared terminal leaflet images. Bars = 2 cm in (B) and 1 mm in (D).

**Figure 6.**
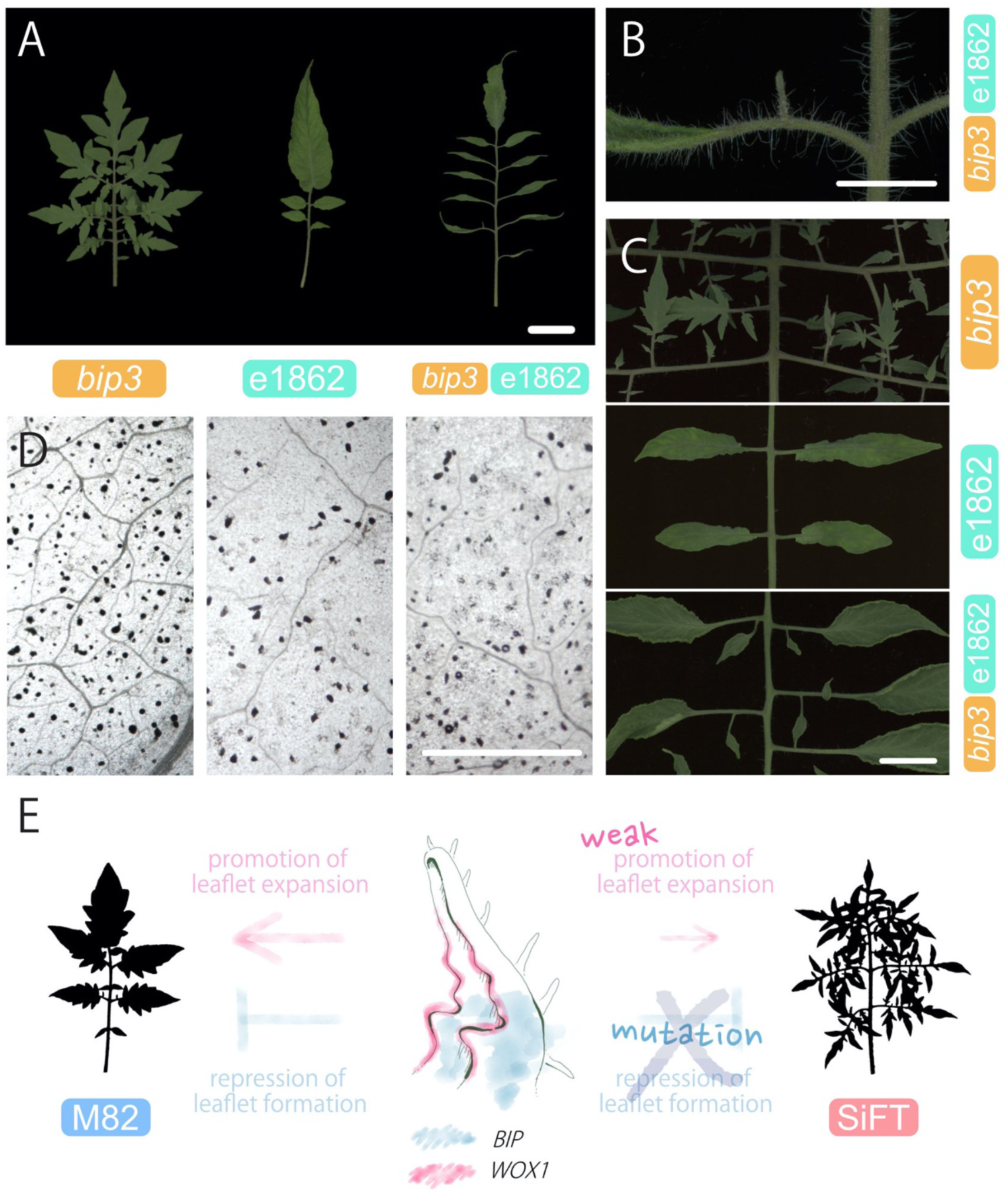
*bip sf* double mutant leaf phenotypes. (A) Mature leaf morphology of *bip3* e1862 double mutant. From left to right: *bip3*, e1862, and *bip3* e1862 double mutant. The 4th leaves were used. (B) Close-up view of a secondary leaflet on a 4th leaf in the double mutant shown in (A). (C) Comparison of secondary leaflets on matured 6th leaf from 60 days old seedlings. (D) Cleared terminal leaflet images of *bip3*, e1862, and *bip3* e1862 double mutant. (E) A schematic model for leaf development in SiFT. Bars = 2 cm in (A) and (C), 1 cm in (B), and 1mm = in (D).

**Figure 7.**
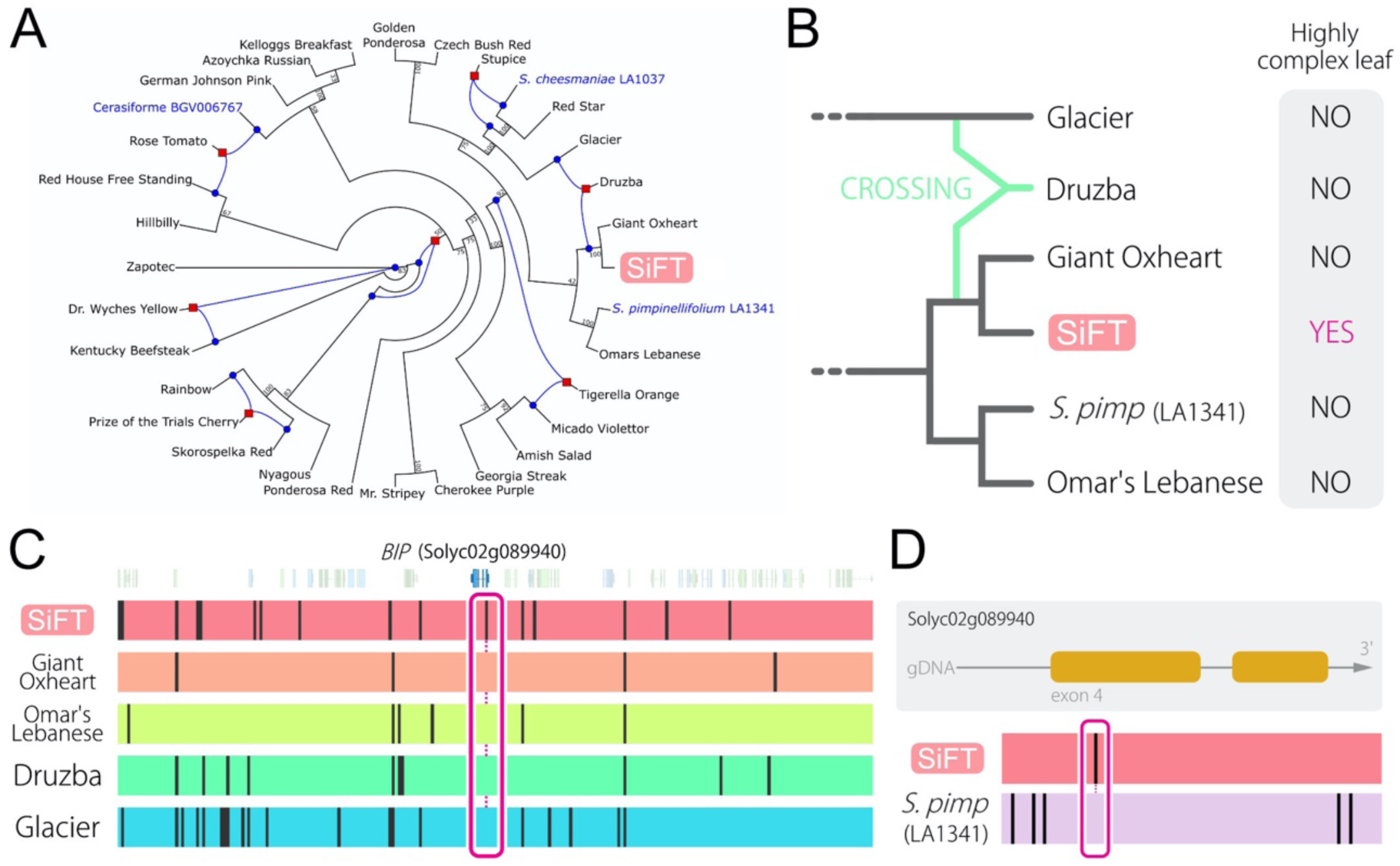
Reconstruction of breeding history and comparison of SNPs data. (A) PhyloNetwork with sequences around the *BIP* locus on chromosome 2. The network describes various biological processes such as hybridization or introgression (Blue lines). The bootstrap values are indicated on branches (only those more than 50% are indicated on the tree). (B) Magnified view of the network shown in (A) focusing on SiFT. Comparison of SNPs data around *BIP* locus (Solyc02g089940) with heirloom tomatoes (C) and a wild species (D). Each vertical black line indicates a SNP.

### Highly complex leaf phenotype seen in SiFT is caused by the *bip* mutation

To verify the effect of the *bip* mutation on leaf phenotypes we investigated the morphology and early development of leaves in *bip3*, a *bip* mutant in the M82 background (12). LC in *bip3* was similar to that of SiFT (Fig. 3A and 3B). Additionally, we confirmed that *bip3* leaf primordia are active in generating multiple leaflets at the P4 stage as seen in SiFT (Fig. 3C). Although the *BIP* gene has been studied in Arabidopsis and tomato, the expression pattern of the *bip* gene in leaf primordia is not known (10, 11). We performed whole-mount *in situ* hybridization and detected *BIP* gene expression in the proximal part of leaf primordium, where leaflet primordia emerge (Fig. 3D). A previous study showed that the expression of *TOAMTO KNOTTED-1* (*Tkn1*), the ortholog of Arabidopsis *KNOX1* gene *BREVIPEDICELLUS*, is increased in the *bip* mutant (10). Quantitative RT-PCR (qPCR) was used to detect elevated level of *Tkn1* expression in SiFT compared to M82 (Fig. 3E). It is known that *KNOX* overexpression increases leaf complexity (12). These data suggest that the highly complex leaf phenotype seen in SiFT is caused by high expression of *Tkn1* facilitated by the *bip* mutation.

Although LC in *bip3* is quite similar to SiFT, the two genotypes have distinctly different leaflet shapes. Deep learning-based nonlinear PCA with leaflet shapes in M82, *bip3*, and SiFT suggested that *bip3* leaf shape is different from that of M82 but not the same as SiFT (Fig. 3F). This trend was confirmed by different methods (Supplementary Fig. 8). Indeed, leaflets of SiFT are narrower than those of bip3 (Fig. 3G). Additionally, LVD in *bip3* is similar to that of M82 and differs from that of SiFT (Fig. 3H and 3I). Thus, the mutation at the *BIP* locus is not sufficient to explain all the leaf phenotypes seen in SiFT.

**Figure 8.**
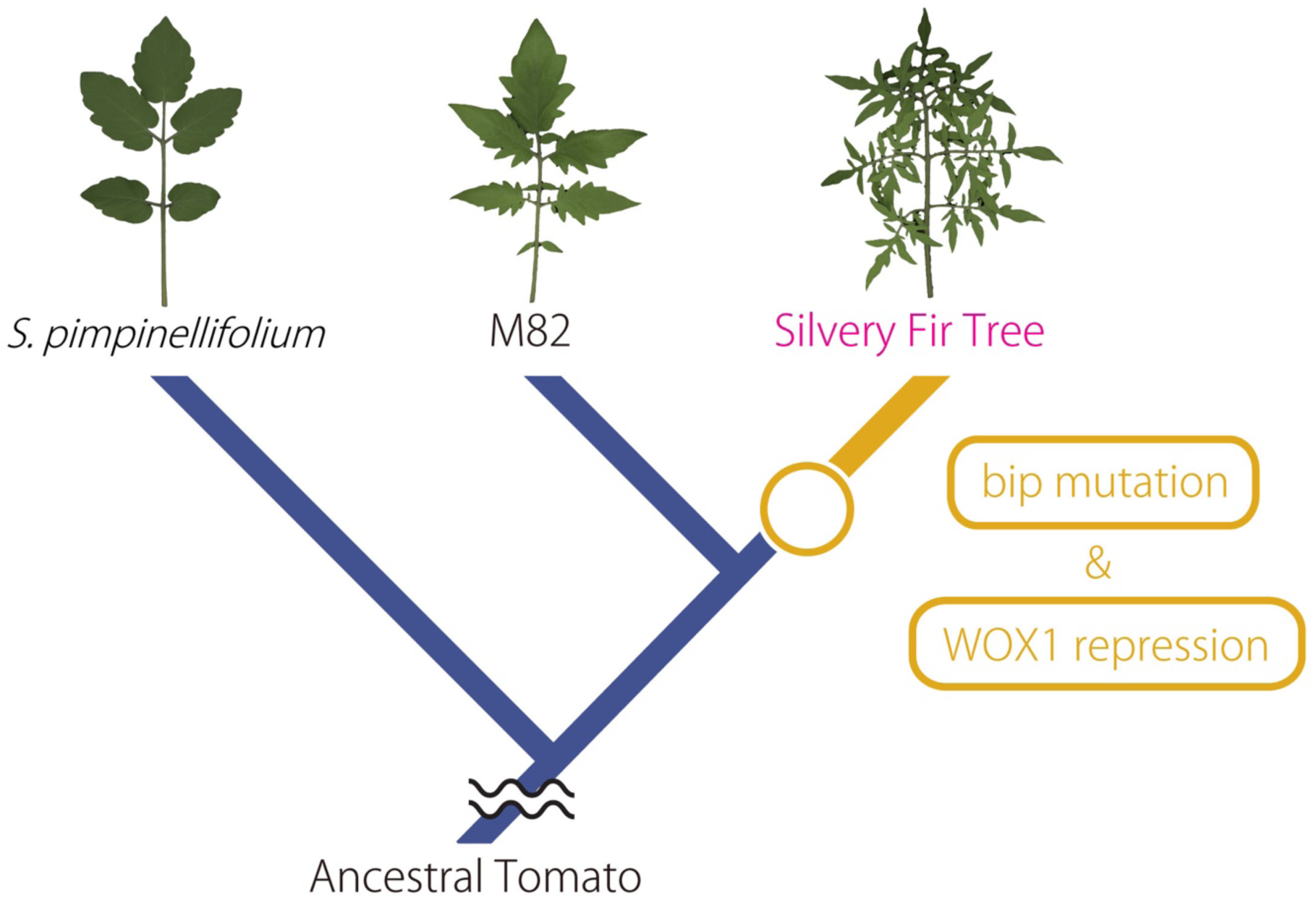
Breeding history and diversification of leaf shape in cultivated Toamto. Simplified phylogeny presented in Fig. 7 with leaf morphologies. Boxes indicate presumed key events during evolution. See text for details.

### Gene co-expression network analysis suggests a role for Sl *WOX1* in regulating leaf phenotypes

To investigate the molecular basis for leaf phenotypes seen in SiFT, we performed RNA-seq and compared differentially expressed genes (DEGs) between M82 and SiFT (Supplementary Table 1). However, the large number of DEGs precluded identification of genes critical in generating differences in leaf shape between two genotypes. Gene co-expression network (GCN) analysis can reveal biologically relevant information to identify molecular mechanisms underlying biological processes (13). Therefore, we constructed GCNs with RNA-seq data of M82 and SiFT and compared these networks to identify key genes responsible for the leaf phenotypes seen in SiFT. The genes used for the network analysis included a set of literature-curated genes involved in leaf development (14). The GCN for M82 showed differences in network structure, edge number, node number, and average of degree between genes when compared to the GCN for SiFT (Fig. 4A and Supplementary Table 2), indicating that many genes involved in leaf development are differentially expressed between the two genotypes. Community structure in the networks was analyzed based on the fast greedy modularity optimization algorithm and GO enrichment analysis by community was performed (14). Two communities (community 1: C1 and community 2: C2) predominate in both networks (Fig. 4A), and GO enrichment analysis by community showed that community 1 is enriched for the same GO terms between M82 and SiFT networks, in particular, GO terms with higher Fold enrichment (>50; full result of the GO enrichment analysis, Supplementary Table 3). On the other hand, community 2 is different between the two networks (Fig. 4B and Supplementary Table 3). In community 2 of the M82 GCN, GO terms such as “cytokinin biosynthetic processes” (GO: 0009691), “cytokinin metabolic process” (GO:0009690), “regulation of cell cycle arrest” (GO: 0071156), and “cellular hormone metabolic process” (GO:0034754; Supplementary Table 3) were more than 100 fold enriched. However, there were no enriched GO terms with high fold enrichment in community 2 of the SiFT GCN (Fig. 4B and Supplementary Table 3), suggesting that genes in community 2 might be crucial for explaining differences in leaf phenotype between M82 and SiFT. To compare the two networks and identify the differences between them, we performed comparative network analysis using the R package “DiffCorr” (15). DiffCorr allows us to find statistically significant differences between two networks. The DiffCorr analysis identified 160 DiffCorr genes, which are differentially correlated genes between two networks (Supplementary Table 4). Those genes should have distinct expression pattern between M82 and SiFT. The 160 DiffCorr genes have distinct expression profiles between M82 and SiFT (Fig. 4C). Additionally, many genes having more differential correlations between two networks show distinct expression patterns, suggesting that those DiffCorr genes are responsible for the difference between the two networks (Fig. 4C). The DiffCorr analysis revealed a *WOX-*like gene (Solyc03g118770) as the most significantly different between the M82 and SiFT GCNs (Supplementary Table 4) and the gene was located in community 2 of the M82 GCN (Supplementary Table 5). Based on phylogenetic analyses and alignments, the *WOX*-like gene is the tomato ortholog of Arabidopsis *WOX1* (Sl *WOX1*; Supplementary Fig. 9 and Supplementary Fig. 10). To understand the role of Sl *WOX1* in leaf development, we focused on the Sl *WOX1* sub-network which consisted of genes showing a direct connection to the Sl *WOX1*. This sub-network showed that the Sl *WOX1* gene is connected to many genes involved in leaf development in M82 GCN (Fig. 4D). Sl *WOX1* expression in SiFT leaf primordia was lower than in M82 samples (Fig. 4E). To check the expression pattern of Sl *WOX1*, we performed whole mount *in situ* hybridization. Sl *WOX1* was expressed at the margins of leaf primordia and leaflet primordia (Figures 4F and 4G). This expression pattern was unaltered in SiFT (Supplementary Fig. 11). *WOX1* is known to express in leaf primordia and is involved in leaf lamina expansion in various plant species (16, 17). Moreover, in the *Medicago truncatula wox1* mutant, leaf vein density was lower than in wildtype (17). Therefore, Sl *WOX1* is likely a candidate gene for controlling both leaf width and leaf vein density in tomato.

### Sl *WOX1* is involved in leaf lamina expansion and leaf vascular development

Sl *WOX1* is located on the long arm of chromosome 3 (Solyc03g118770; https://solgenomics.net/feature/17777644/details) and previous studies in several plant species indicate that the *wox1* mutants have narrower leaflets and low LVD (16, 17). Therefore, we mutated Sl *WOX1* in M82 plants using the CRISPR/Cas9 system and obtained a null mutation, referred to here as CR-*wox1-1* (Fig. 5A). CR-*wox1-1* plants showed narrower leaves (Fig. 5B and 5C) and lower LVD than M82 (Fig. 5D), matching the phenotype described in *wox1* mutants in other species (16, 17). We noticed that these phenotypes are quite similar to those of a classical tomato mutant, *solanifolia* (*sf*). *sf* is known to have narrower leaflets and reduced vascular density (18) and our rough mapping of the *solanifolia* mutation had identified a genomic location close to that known for the *WOX1* locus (at the end of chromosome 3; Supplementary Fig. 12). We obtained the two known alleles of the mutation (*sf* and *sf^wl*) and another allele (e1862) from the Tomato Genetics Resource Center (TGRC; http://tgrc.ucdavis.edu) and Genes that makes Tomato (http://zamir.sgn.cornell.edu/mutants/), respectively. *sf, sf^wl*, and e1862 arose in the Pearson, ROMA, and M82 backgrounds, respectively. *sf* has a 1 bp substitution (G to A) at position 230 in Sl *WOX1* resulting in an amino acid swap from arginine to histidine in the conserved homeodomain (Fig. 5A and Supplementary Fig. 13). *sf^wl* and e1862 have a 1 bp substitution (G to A) at a splice site between intron 3 and exon 4, which results in 10 bp shift to the next splice site (Supplementary Fig. 13 and Supplementary Fig. 14). As a result, these mutants have a 10 bp deletion from position 595 to 604 resulting in a premature stop codon, which truncates the WOX1 protein such that it lacks the conserved WOX domain (Fig. 5A and Supplementary Fig. 13 and Supplementary Fig. 14). All these mutants showed narrower leaflets and low LVD compared to their background genotypes (Fig. 5B to 5D). Hence, phenotypes seen in these classical mutants were same as that of CR-*wox1-1*. Hereafter, we refer to the Sl *WOX1* gene as *SOLANIFOLIA* (*SF*). These results indicated that *SF* functions in leaflet outgrowth and vascular development in tomato leaves, performing a role similar to that of *WOX1* in Arabidopsis and Medicago (16, 17, 19). These results confirm the role of reduced expression of *SF* in conferring narrower leaflet and lower LVD phenotypes in SiFT. Promoter sequences of *SF* in SiFT show no SNPs compared to the reference tomato genome (Supplementary Fig. 15), coincident with the fact that the pattern of expression *SF* is unaltered in SiFT compared to M82 (Supplementary Fig. 11). A previous study suggested that *ARF3* suppresses *WOX1* expression in Arabidopsis (20). In SiFT, expression level of the *ARF3* ortholog (Solyc02g077560) was higher than that seen in M82 (Supplementary Fig. 16A) and the *ARF3* ortholog is known to be expressed in leaf primordia (21). Additionally, the SiFT *ARF3* promoter has a SNP that generated a new ZF-HD motif, which is known to be involved in binding of HB33 transcription factor (22) (Supplementary Fig. 16B). This *HB33* ortholog in Toamto (Solyc04g080490) is expressed in leaf primordia (Supplementary Fig. 17). Therefore, this gain-of-function SNP might explain the alteration in *SF* repression in SiFT.

### *bip sf* double mutant shows highly complex, narrow leaves, and low LVD

Since SiFT has a *bip* mutation and *SF* repression, we generated a *bip sf* double mutant to investigate leaf phenotypes. Leaves in double mutant between *bip3* and e1862 had more leaflets than those of e1862 and the leaflets were narrower than those of *bip3* (Fig. 6A). Sometimes, secondary leaflets were observed on the 4th leaf (Fig. 6B) and the secondary leaflets became more obvious in higher order leaves in the double mutant (Fig. 6C). Additionally, LVD in the double mutant was lower than that in *bip3* (Fig. 6D), suggesting that these phenotypes are additive. Moreover, these trends were confirmed by another double mutant with a different combination of mutants (Supplemental Fig. 18). However, leaf morphology in the double mutant was not exactly the same as that of SiFT. Leaflets in leaves of SiFT have many robes (Fig 1B), leaves of the double mutant, however, do not have obvious lobes (Fig. 6A and Supplemental Fig. 18). This may be due to the difference between reduced expression and complete loss of function of the *SF* in the two genotypes. Indeed, *sf* single mutants having truncated WOX1 protein such as CR-*wox1-1, sf^wl*, and e1862 do not have any lobes on leaves, whereas a weaker phenotype mutant, *sf*, shows lobed leaves (Fig. 5B). These results suggest that a mutation at *bip* and *WOX1* repression lead to highly complex and narrower leaves with reduced leaf vein density in SiFT, respectively (Fig. 6E).

### Phylogenetic placement of SiFT and the *bip* mutation

Several mutations at the *BIP* locus have been described (10). However, the *BIP* mutation in the SiFT genome is different from these. Although previous studies constructed phylogenies with heirloom tomatoes (23), they were generated with whole-genome sequencing data. In order to understand the history of the *bip* mutation on chromosome 2, we first constructed a phylogenetic tree based on whole-genome sequencing data from 106 heirloom tomatoes (http://www.tomatogenome.net/accessions.html) to know the rough relationship among them (Supplemental Fig. 19). Subsequently, we constructed a phylogenetic network using the “PhyloNetworks” package in Julia (24) to estimate whether introgressions from other tomatoes occurred in SiFT or not. This package allows us to discern various biological processes such as hybridization, introgression, or horizontal gene transfer. We used sequences around the *BIP* locus on chromosome 2 from 32 representative tomatoes based on the phylogeny with 106 heirloom tomatoes. *S. pimpinellifolium*, which is thought to be the progenitor wild species for domesticated tomato, was used as an outgroup in the heirloom tomato phylogeny. *S. cheesemaniae* and *S. lycopersicum* var. *cerasiforme* were also used (Fig. 7A). The network indicated that a US heirloom tomato, Giant Oxheart (GiO), is sister to SiFT, however GiO does not show the highly complex leaf phenotype characteristic of SiFT and lacks the mutation in *BIP* (Fig. 7B and 7C). The phylogeny suggests that Druzba is the result of a cross between an ancestor of Glacier and an ancestor of Giant Oxheart/SFT. However, the *bip* mutation seen in SiFT does not exist in Druzba either (Fig. 7B and 7C). Additionally, the wild species, *S. pimpinellifolium* (Fig. 7D), and the other tomato varieties do not harbor the SiFT specific mutation at *BIP* (Supplemental Fig. 19 and Supplementary Table 6). These data suggest that the *bip* mutation in SiFT is likely a *de novo* mutation, instead of standing genetic variation.

## Discussion

We found that SiFT, an heirloom tomato, has a highly complex leaf phenotype and carries a mutation in the *BIP* gene, which encodes a *BEL-LIKE HOMEODOMAIN* protein (Solyc02g089940). Leaf complexity in the *bip3* mutant was remarkably similar to that of SiFT. *BIP* is expressed at the proximal end of developing leaf primordia, where leaflet primordia emerge. Previous studies demonstrated that *KNOXI* genes are overexpressed in leaves of *bip* mutants (10, 11) and the overexpression of *KNOX1* gene in leaves dramatically increases leaf complexity by prolonging specific stages of leaf development (25). Indeed, *Tkn1*, a tomato *KNOX1* gene, is highly expressed in SiFT leaf primordia and SiFT and *bip3* leaf primordia exhibit prolonged morphogenesis. In Arabidopsis, *saw1 saw2* double mutant showing ectopic *KNOX1* expression has increased leaf serrations (10, 11), indicating that *BLH* genes including *BIP, SAW1*, and *SAW2* act to limit leaf margin growth and the function appears to be conserved between them. Therefore, the *bip* mutation found in SiFT is the likely cause of increasing in *Tkn1* expression, prolonging morphogenesis, and increasing complexity in SiFT leaf primordia. We also found that leaflet shape in SiFT is narrower than *bip3*, which is confirmed by deep learning-based nonlinear PCA and leaf shape analysis. Additionally, leaf vein density in SiFT was lower than in *bip3*. To identify the genetic alterations beyond *BIP* that explain the rest of leaf phenotypes seen in SiFT, we used comparative gene co-expression network analysis. The *WOX1* ortholog (Solyc03g118770) had the most altered correlations between the M82 and SiFT co-expression networks. The expression level of the *WOX1* ortholog in SiFT is lower than M82. Additionally, *wox1* mutants in Arabidopsis and Medicago are known to have narrower leaves compared to WT (16, 17). A CRISPR/Cas9 *wox1* mutant in Tomato showed narrower leaves and lower vascular density compared to WT. Moreover, we found that a classical tomato mutant, *solanifolia (sf)*, harbored a deleterious mutation in the *WOX1* ortholog and those *sf* mutants showed narrower leaves and lower vascular density. Whole-mount in situ hybridization demonstrated that, similar to Arabidopsis and Medicago (16, 17), *SF* is also expressed at the margin of tomato leaf and leaflet primordia, consistent with the phenotype of leaf margins in *sf* mutants. In Arabidopsis, *WOX* genes promote lamina outgrowth through regulation of cell proliferation in cells expressing *WOX1* and their surrounding cells (16). Since leaf vascular development is influenced by this marginal blade outgrowth (26), we propose that *SF* functions in leaf lamina outgrowth and couples this growth feature with vascular patterning. The causal mutation that leads to *SF* repression in SiFT might be a SNP in ARF3 promoter region in the SiFT genome, however this needs further analysis. *WOX1* is present in the early-diverging angiosperm, *Amborella trichopoda* (27), but the ancestral function in angiosperms is still unknown. Additionally, no *WOX1* homologs have been identified in monocots (17, 28). These facts suggest that the function of this *WOX1* gene in leaf development appears to be conserved at least across the eudicots (e.g. Arabidopsis, Medicago, and Tomato).

Leaves of *bip* and *sf* double mutants are more complex than those of *wox1* and narrower than those of *bip3*. Moreover, the double mutant showed low LVD. A recent study suggested that the regulation of local growth and differentiation in leaf primordia leads to diversity in leaf shape (29). *BIP* and *SF* are thought to regulate local growth and differentiation in leaf primordia: *BIP* functions in the proximal part of leaf primordia and *SF* functions at the marginal part. Therefore we conclude that the highly complex and narrower leaf with reduced leaf vein density seen in SiFT is caused by a combination of a mutation at *bip* and another as yet unknown second site mutation that leads to *SF* repression (Fig. 8).

A phylogenetic tree constructed with WGS data and a phylogenetic network constructed with sequences around *BIP* locus revealed that German Red Strawberry and Giant Oxheart are sister to SiFT, respectively. However, they lack the *BIP* mutation and have regular leaf shape. Moreover, none of the other varieties or wild species harbor the same mutation at *BIP* seen in SiFT. Although a wild tomato species *S. galapagense* has increased leaf complexity, the increased leaf phenotype is linked to promoter alterations in an atypical *KNOX1* gene *PETROSELENUM*, not in *BIP* (10). Therefore, these data indicate that the *BIP* mutation seen in SiFT is a *de novo* mutation that occurred during breeding and is not likely to be an introgression from other varieties or wild species.

These are consistent with the fact that there is no cultivated tomato showing SiFT-like leaf shape. This uniqueness of leaf shape in SiFT is achieved by the combination of a mutation at *bip* and *SF* repression, leading to use of this variety as an ornamental and landscaping plant (6). Emerging data suggest that leaflet shape affects fruit sugar content in tomato (30, 31). Therefore, identification of these novel mutations not only provides new insights into the breeding history of heirloom tomatoes, but also suggests potential targets for enhancing sugar content to improve fruit quality in tomato.

## Methods

### Plant Materials

The following lines and mutants were provided by the Tomato Genetics Resource Center, University of California, Davis (USA; https://tgrc.ucdavis.edu/): *S. lycopersicum* cv M82 (LA3475), *sf*^wl (LA2012), *sf* (LA2311), and Pearson (LA0012). SiFT and ROMA were from our own stocks. *bip3* (e1444m2) was obtained from the saturated mutation library of tomato (32). e1862 was obtained from the Genes that makes Tomato (Israe<u>l; http://zamir.sgn.cornell.edu/mutants/).</u>

### Growth conditions

Tomato seeds were soaked in 50% bleach for 10 min, rinsed 3 times with water, and placed on water dampened paper towel in Phytatrays (Sigma Aldrich). Seeds were incubated in the dark at room temperature for 3 days then transferred to a growth chamber set at 22°C under long-day conditions (16 h light; 8 h dark) for 4 days. After approximately 7 days, seedlings had expanded cotyledons. These were then transplanted to 24-cell seedling propagation trays and grown in the chamber for a 35 days as described previously (33) and arranged in a randomized block design. The shoots or leaf primordia were frozen in liquid nitrogen just after sampling, and then stored at –80°C until use for DNA and RNA extractions. All F2 plants were grown in a field in the University of California, Davis with an interplant spacing of 30 x 30 cm^2^ for transplanting.

### Morphological Observations

For morphological observations and collecting tissues for RNA extractions, shoots and leaves were dissected under the dissection microscopes (Discovery.V12; ZEISS). To determine the vascular density of leaves, the 6th leaves were used and cleared using an ethanol and 50 % bleach following Rowland *et al*. (2019). The samples were then photographed under the microscopes (ECLIPSE E600; Nikon), and vascular length per unit area was determined using ImageJ software (http://rsb.info.nih.gov/ij/) (n = 4). Leaf complexity was determined by counting the numbers of leaflets and intercalary leaflets on a fully developed leaf (n = 29). Traditional leaf shape analysis was performed following (34). Leaf complexity and leaflet shapes were analyzed for leaves collected from the chamber. The leaf complexity measures included all leaflets present on the leaf. After complexity was obtained the primary leaflets were removed and used for imaging and analysis of shape and size. The intercalary and secondary/tertiary leaflets were not used for shape analysis due to their smaller size and irregular shapes. The binary images were then processed in R using MOMOCS, a shape analysis package (35).

### Phylogenetic Analyses of Isolated Genes

The predicted amino acid sequences of isolated genes were aligned using ClustalW and readjusted manually. Phylogenetic trees were reconstructed using MEGA6 (36) using the neighbor-joining method (37) (38). Bootstrap values were derived from 1000 replicate runs. The ML phylogenetic tree with the highest log likelihood is shown. Initial trees for the heuristic search were obtained automatically: Neighbor-Join and BioNJ algorithms were applied to a matrix of pairwise distances estimated with MCL, and then the topology with a superior log likelihood value was selected. The tree is drawn to scale, with branch lengths measured in the number of substitutions per site.

### Whole-mount In Situ Hybridization

Portions of genes isolated in pCR 2.1 (Invitrogen) were amplified by PCR using the universal primers M13_F (−20) (GTAAAACGACGGCCAC) and M13_R (CAGGAAACAGCTATGAG). The amplified fragments were then used to produce digoxigenin (DIG)-labeled sense and antisense RNA probes using a DIG RNA Labeling Kit (Roche). Whole-mount in situ hybridization was performed following (39). Shoots were fixed in 1x PBST containing 4% (w/v) paraformaldehyde, 1% (w/v) glutaraldehyde. Fixed samples were dehydrated in an ethanol series. The dehydrated samples were stored in 100% methanol at –20°C until use for the experiment. DIG-labeled sense and antisense RNA probes were synthesized with T7 RNA polymerase (Roche). For immunological detection, the samples were incubated in detection buffer containing NBT-BCIP (Roche) at 25 °C for several hours or 4°C overnight. Photographs were taken using an ECLIPSE E600 (Nikon). The experiments were performed at least three times.

### Quantitative Real-Time PCR

Total RNA was extracted from leaf primordia of plants grown for a month and used to synthesize cDNA, as described above. The quantitative RT-PCR analysis was conducted using the following gene-specific primer pairs: Tkn1_RT_F and Tkn1_RT_R; SlWOX1_RT_F and SlWOX1_RT_R; and SlGAPDH_RT_F and SlGAPDH_RT_R (Supplemental Table 7). Real-time PCR amplification was performed using the iTaq Universal SYBR (BIO-RAD) in a iQ5 Real-Time PCR Detection System (BIO-RAD). Experiments were performed in triplicate from three independent tissue RNA extractions. Expression was normalized to the Sl *GAPDH* control.

### Deep learning-based nonlinear PCA

For nonlinear PCA on image data, we used leaflet images from M82, *bip3*, and SiFT for the analysis (4th leaf; N < 55) and adopted a pre-trained neural network with the ImageNet dataset, VGG19 (40), as feature extractor. Instead of the original scanned images, binary silhouette images were fed into the network in order to avoid the effects of non-morphogenetic features such as leaf color. We extracted images features from an intermediate layer, “block4_pool” through Keras 2.3.1 library (https://keras.io). Then the linear PCA was applied on the image features. We performed no training of the neural network with our data, so that the feature extraction was completely agnostic on which genotypes the leaves came from.

### DNA-Seq and RNA-Seq Library Preparation and Sequencing

DNA-Seq libraries for BSA were prepared following (41). DNA was extracted using GeneJET Plant Genomic DNA Purification Mini Kit (Thermo Scientific, Waltham, MA, USA) from plants grown for a month. DNA-Seq libraries for phylogenic analysis were prepared based on BrAD-seq (42) with the following modifications: After DNA fragmentation with Covaris E220 (Covaris, Inc. Woburn, MA, USA), the fragmented DNA was end-repaired, A-tailed, and adapter ligated with Y-adapter. Enrichment PCR was then performed with the adapter ligated product as described Townsley et al., 2015. After final library cleanup with AMPure beads (Beckman Coulter, Brea, CA, USA), RNA-Seq libraries were prepared following (42) from four biological replicates of proximal and distal regions of leaf primordia at four developmental stages (meristem + P1-P3; P4; P5). DNA-Seq libraries were sequenced at Novogene (Novogene Inc. Sacramento, CA, USA). RNA-Seq libraries were sequenced at the University of California Berkeley Vincent J. Coates Genomics Sequencing Laboratory using the HiSeq 2000 platform (Illumina Inc. San Diego, CA, USA).

### SNP calling and Allele frequency analysis with DNA-seq data and Phylogenetic analysis with Whole genome sequencing data

To detect SNPs in SiFT genome and perform phylogenetic analysis, all variants detected by CLC Genomics Workbench 11.0 (CLC Bio, a QIAGEN Company, Aarhus, Denmark). After read mapping and local realignment, Fixed Ploidy Variant Detection function was used for calculation of allele frequency. For phylogenetic analysis, the data were exported as vcf files. The SNPRelate package for R (43) was used to determine the variant positions that overlapped between cultivars and then all sequences combined into a single gds file. This file was run through SNPhylo (44) with the following parameters: The linkage disequilibrium was set to 1.0, as we wanted to exclude as few variants as possible based on this factor, minor allele frequency was set to 0.05, and the Missing rate was set to 0.1. One thousand bootstraps were performed for confidence intervals and significance. *S. pimpinellifolium* was used as the out group. The output bootstrapped tree was displayed in MEGA6 (36).

### Mapping, Normalization, and Network analysis with RNA-Seq data

The 50 bp single-end sequence reads obtained were quality trimmed and parsed to individual libraries using custom Perl scripts. All reads were mapped to the ITAG2.4 genome build (downloadable from http://solgenomics.net/itag/release/2.4/list_files) using RSEM/eXpress with the default parameters (45). The uniquely mapped read data was normalized using the Bioconductor package EdgeR ver. 2.11 with the trimmed mean of M-values method. Bioinformatics and statistical analyses were performed on the iPLANT (Cyverse) Atmosphere cloud server (46). Gene Co-expression network analysis was performed following (14) by using the R script. The R script for RNA-Seq gene coexpression network analysis deposited on GitHub (Link: https://github.com/Hokuto-GH/gene-coexpression-network-script). For GO enrichment analysis, we used GENEONTOLOGY enrichment analysis tools (http://geneontology.org/docs/go-enrichment-analysis/). DiffCorr analysis was performed following (15). The normalized count data from M82 and SiFT was used for the analysis. DiffCorr genes were then analyzed to identify the most different gene between two genotypes at a 0.005 FDR cut-off. To analyze and visualize the DiffCorr genes, Cytoscape was used (https://cytoscape.org/). The number of Edges of each DiffCorr gene was calculated by analyze network function in the Cytoscape. Then, the numbers were compared to figure out the most different gene between two genotypes.

### CRISPR–Cas9 mutagenesis and plant transformation

CRISPR–Cas9 mutagenesis and generation of transgenic plants was performed following REF. Guide (g) RNAs for *SF/SlWOX1* (Solyc03g118770) were designed using the CCtop (https://crispr.cos.uni-heidelberg.de/help.html) and two gRNAs were designed (Fig. 5). Vectors were assembled using the Golden Gate cloning system as described (47). Final binary vector was transformed into the tomato cultivars M82 by *Agrobacterium tumefaciens*-mediated transformation. The transformation was performed at the Ralph M. Parsons Foundation Plant Transformation Facility (University of California, Davis). The first-generation (T_0_) transgenics were genotyped using GT-seq following (48). It revealed a single nucleotide substitution (C to A) in gRNA2 (g2) region. Unfortunately, there were no T_0_ transgenics having mutation in the region of gRNA1 (g1) region. After the genotyping and self-pollination in the green house, we obtained T_1_ plants having mutated *sf/slwox1* gene. First, we screened those plants by leaf phenotypes because *wox1* mutants must have narrower leaflets compared to WT based on previous studies with various kinds of plant species. Then we did genotyping by sequencing to confirm whether each individual has the *sf/slwox1* mutation or not.

### PhyloNetwork Analyses

To perform phylogenetic analysis, all SNPs detected by CLC Genomics Workbench 11.0 (CLC Bio, a QIAGEN Company, Aarhus, Denmark) from whole genome sequencing obtained from the 360 genomes project (49) were exported as a vcf file. The VCFtools package (50) was to convert vcf files to fasta files and these sequences were aligned using ClustalW. All aligned SNPs from the two megabase region surrounding the BIP gene for 32 cultivars were run through the TICR pipeline (51). They were then analyzed using PhyloNetworks with default settings with the following exceptions: the number of runs was set to 10 and Nfail was set to 10. After the hybrid network was obtained bootstrap analysis was done in PhyloNetworks using default settings with the following exceptions: Runs was set to 10 and Nfail was set to 10. These adjustments were made to decrease processing time. The bootstrapped tree was output in Dendroscope (52).

### Statistical analysis

All statistical analyses were performed using JMP (JMP Pro 14.0.0, 2018) software. To determine statistical significance, measurements were modeled using general linear regression model and tested by a one-way ANOVA followed by Tukey’s honestly significant difference, if necessary.

## Data availability

All data is available in the main text or the supplementary materials. All DNA-Seq and RNA-Seq raw data are deposited on DDBJ DRA009167-009182 (BioProject: PRJDB8552). Source Data files for all main and supplementary figures are available in the online version of the paper. All additional data sets are available from the corresponding author on request.

## Acknowledgements

This work was supported by grants from the USDA NIFA (2014-67013-21700), NSF; IOS (1558900), and JSPS KAKENHI (19K23742 and 20K06682 to H.N. and 19H05670 to Y.K.). Hokuto Nakayama was a recipient of a JSPS Fellowship (13J00161). We are grateful to Lauren Hughes for generating the initial cross between SiFT and M82, Donnelly West for the F2 seeds, and Amber Flores for helping with field measurements.

## Author contributions

H.N. initiated the project. N.R.S. supervised the project. H.N. and N.R.S. designed experiments. H.N. performed the majority of the experiments and analyses and prepared figures. S.D.R., Z.C., K.Z., J.C., and Y.K. performed experiments. H.N. and S.D.R. analyzed the data. H.N. and N.R.S. wrote the paper with the input from all authors.

## Corresponding Author

Correspondence to Neelima R. Sinha

