## Supplemental Figures for "Leaf form diversification in an heirloom tomato results from alterations in two different *HOMEOBOX* genes": Supplementary_figures.pdf

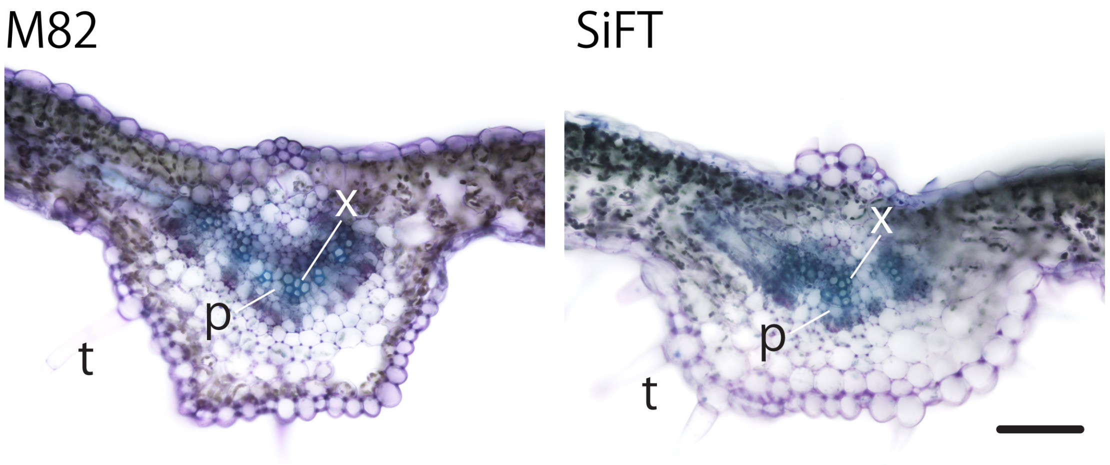


**Supplementary figure 1.** Leaf anatomy in M82 and SiFT.

Transverse sections of 4th leaves in M82 and SiFT. p, phloem; t, trichome; x, xylem. Top of the image is the adaxial side. Bar = 100 $\mu$m.

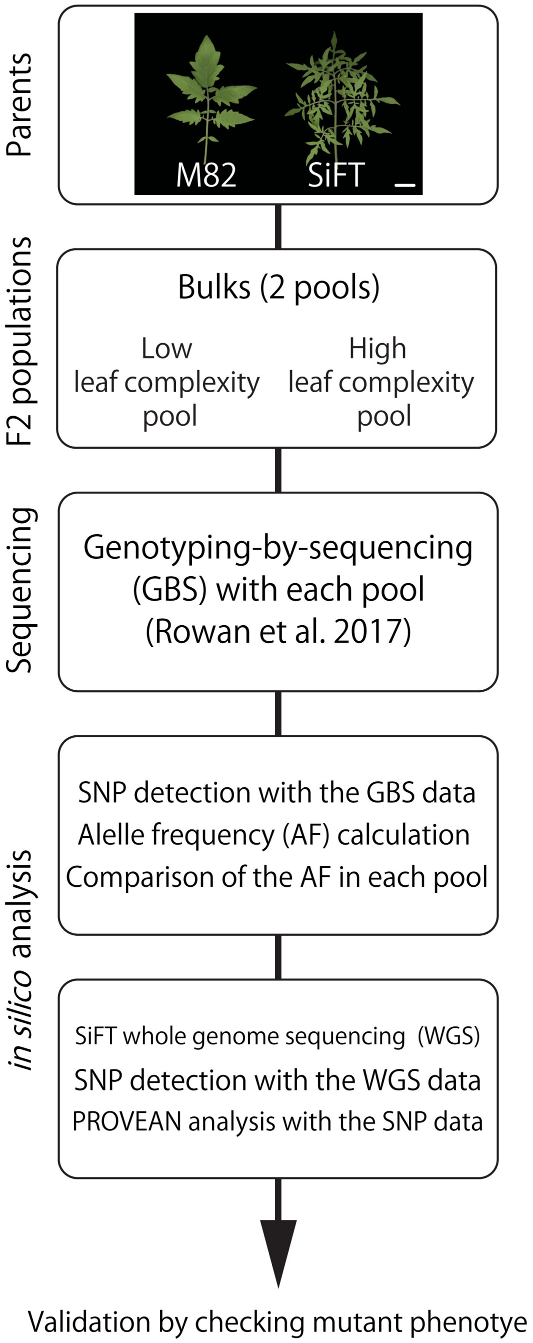


**Supplementary figure 2.** Overview of combination in BSA, WGS, and PROVEAN to detect genes involved in the regulation of leaf complexity in this study.

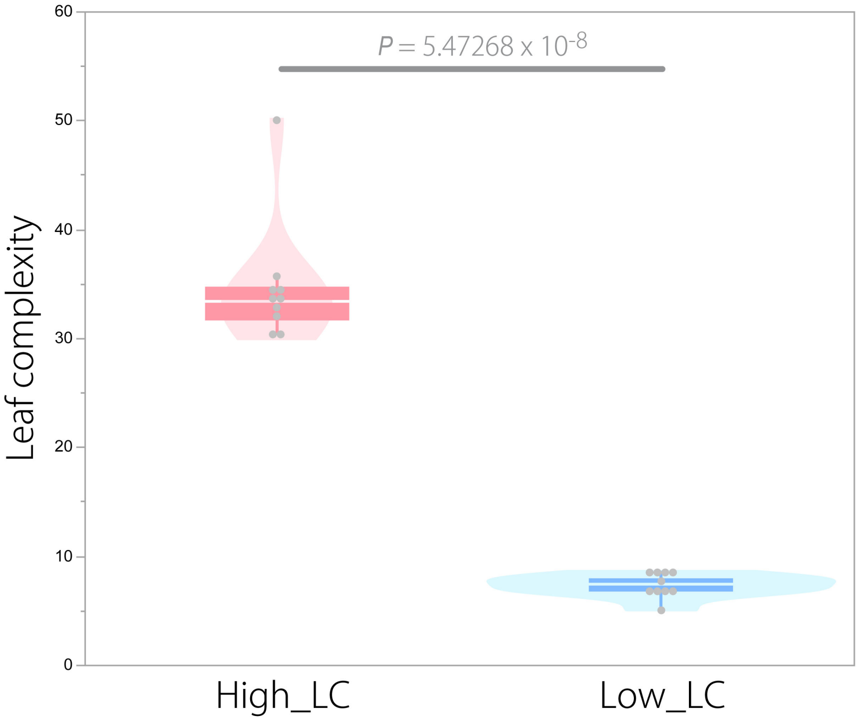


**Supplementary figure 3.** Comparison of leaf complexity between two bulks. Comparison of leaf complexity (LC) between High_LC bulk and Low_LC bulk. (n = 10). $p = 0.0000000547268$ (Welch's *t*-test).

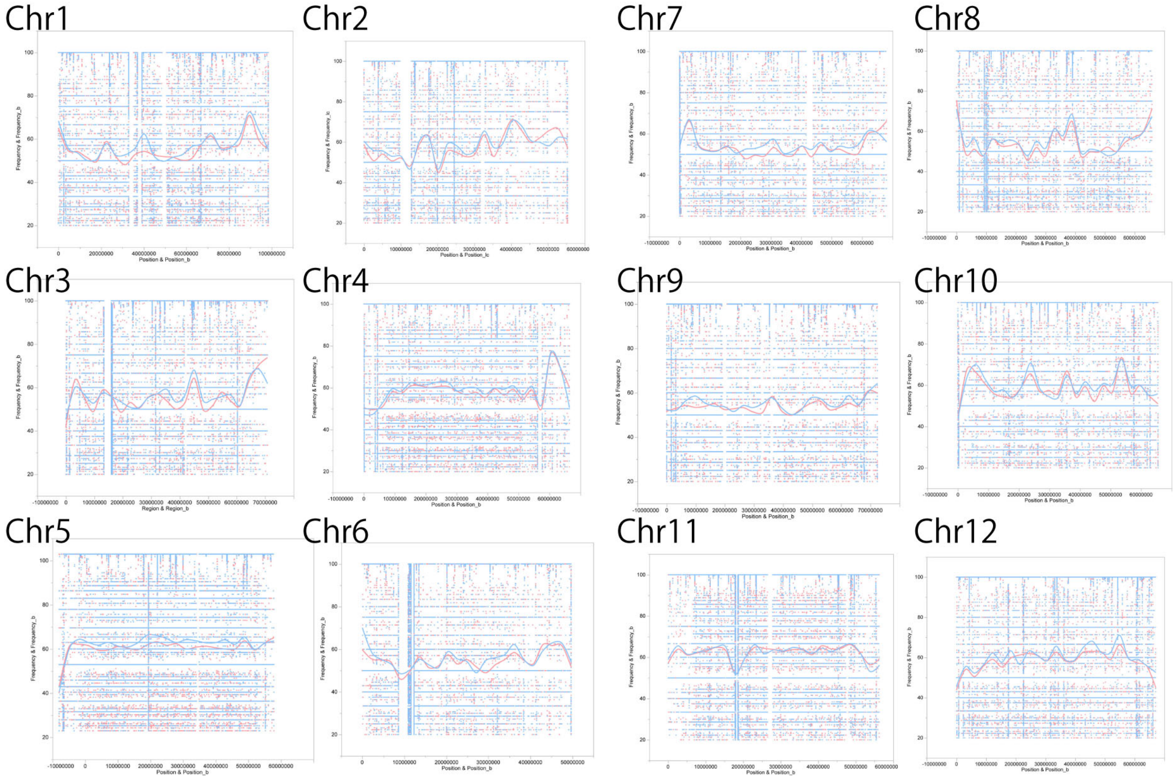


**Supplementary figure 4.** Comparison of allele frequency between two bulks in all chromosomes.

Allele frequency between different pools of segregating populations (red: high complexity pool; blue: high complexity pool) is shown.

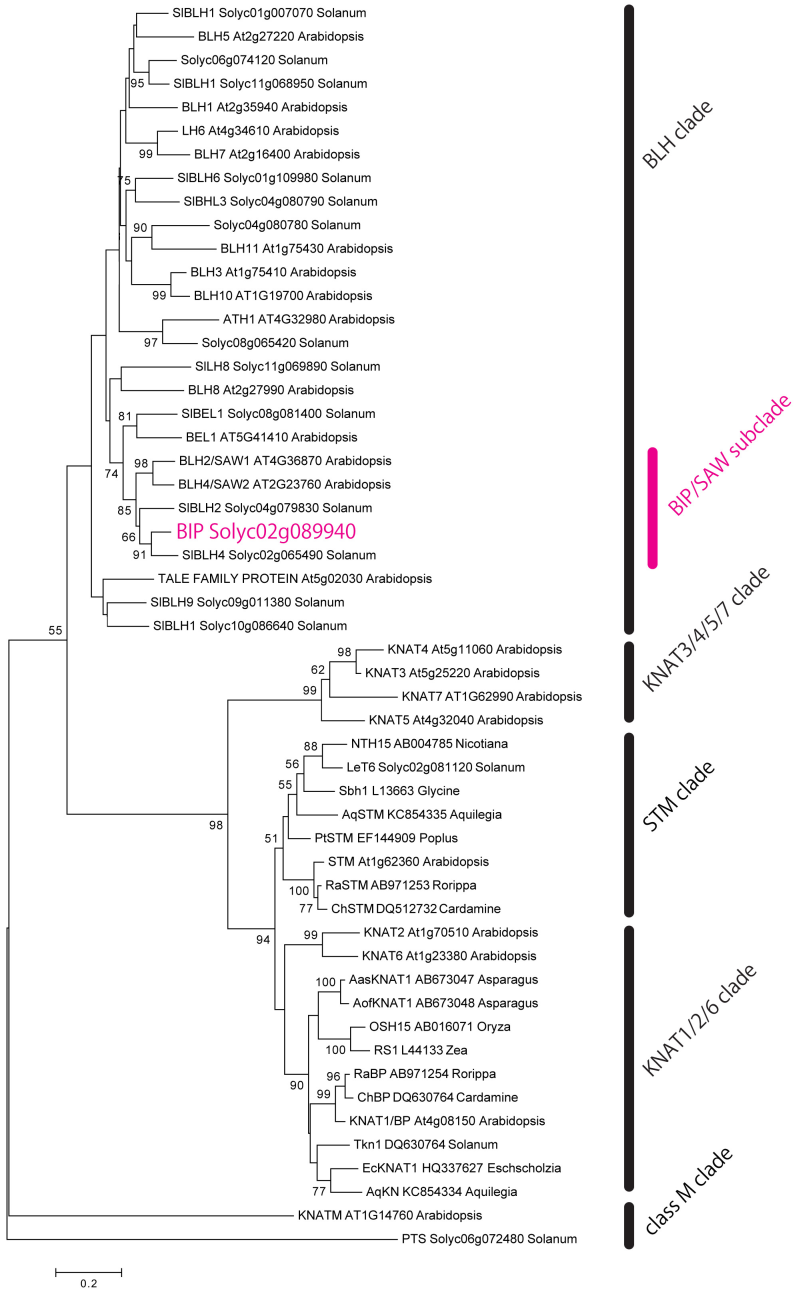


**Supplementary figure 5.** A phylogenetic tree of *BELL/KNOX* orthologs.

A phylogenetic tree of *BELL/KNOX* orthologs resulting from Neighbor-joining method by MEGA. *BIP* is located in a *BLH* clade in the phylogeny. The bootstrap values are indicated on branches (only those more than 50% are indicated on the tree).

```
SFT        MGIATQPITLRPCHKYPVIQQVVENLNSMSQDYHHH--------SLFSFPNGFERSQAEQ    52
M82        MGIATQPITLRPCHKYPVIQQVVENLNSMSQDYHHH--------SLFSFPNGFERSQAEQ    52
SAW1       MGITKT-------SPNTTILLKTFHNNSMSQDYHHHHHHNQHQGGIFNFSNGFDRSDSPN    53
SAW2       MGL------------------ATTTSSMSQDYHHH-------QGIFSFSNGFHRSSSTT    34
           **:                   .   .*********        .:*.* ***.**.:

           QQHQHQQAQQIRRDKLRVQGFEPFEDETSGLPTVYETAGMLSEMFNFPPGNAATNAAELL    112
           QQHQHQQAQQIRRDKLRVQGFEPFEDETSGLPTVYETAGMLSEMFNFPPGNAATNAAELL    112
           LTTQQKQEHQR--V-----EMDEESSVAGGRIPVYESAGMLSEMFNFPGSSGGGRDLDLG    106
           ---------HQ--E-----EVDESAVVSGAQIPVYETAGMLSEMFAYPGGGGGGSGGEIL    78
                    :          .:     :..   ***:******** :* ....    ::

           ETQFNPNFRQPNPRIHAAAAMGNEWFGNHRQGMVVGGGASSQPLGYAKNHTDSMQLFLMN    172
           ETQFNPNFRQPNPRIHAAAAMGNEWFGNHRQGMVVGGGASSQPLGYAKNHTDSMQLFLMN    172
           QS-FRSNRQL----L-----------EEQHQNIP-----AMNATDSATATAAAMQLFLMN    145
           DQ---STKQL----L-----------EQQN------------------------------    90
           :     . :     :            ::.

           PQPRSPSPSPPN-------STSSTLHMLLPNPSSTPTLQ-GFPNP------AEGSFGQFM    218
           PQPRSPSPSPPN-------STSSTLHMLLPNPSSTPTLQ-GFPNP------AEGSFGQFM    218
           PPPPQQPPSPSSTTSPRSHHNSSTLHMLLPSPSTNTTHHQNYTNHMSMHQLPHQHHQQIS    205
           ---------------RHNNNNNSTLHMLLPNHHQGFA----FTDENTM----QPQQQQHF    127
                               ..********.     :    : :        .    *

           TWGNGGASAATATHHLNAQNEIGGVNVV-----ESQGLSLSLSSSLQHKAEELQMSGEAG    273
           TWGNGGASAATATHHLNAQNEIGGVNVV-----ESQGLSLSLSSSLQHKAEELQMSGEAG    273
           TWQSSP---DHHHHHHNSQTEIGTVHVENSGGHGGQGLSLSLSSSLEAAAKAEEYRN---    259
           TWPSSS---SDH---HQNRDMIGTVHVE-----GGKGLSLSLSSSLA-AAKAEEYRS---    172
           ** ..           : :  ** *:*       .:**********   *:  :  .

           GMLFFNQG-G-------SSTSGQ-YRYKNMNMGGSGISPNIHQVH----------VGYGS    314
           GMLFFNQG-G-------SSTSGQ-YRYKNMNMGGSGISPNIHQVH----------VGYGS    314
           --IYYGANSS------NASPHHQYNQFKTLLANS---SQHHHQVLNQFR---SSPAASSS    305
           --IYCAAVDGTSSSSNASAHHHQFNQFKNLLLENSSSQHHHHQVVGHFGSSSSSPMAASS    230
             ::     .       ::   *  ::*.:   .   . : ***            . .*
                       SR/KY domain
           SLGVVNVLRNSKYAKAAQELLEEFCSVGRGKLKKNNNKAAANNPS--GG------ANNEA    366
           SLGVVNVLRNSKYAKAAQELLEEFCSVGRGKLKKNNNKAAANNPS--GG------ANNEA    366
           SMAAVNILRNSRYTTAAQELLEEFCSVGRGFLKKNKLG-NSSNPNTCGGDGGGSSPSSAG    364
           SIGGIYTLRNSKYTKPAQELLEEFCSVGRGHFKKNKLSRNNSNPNTTGGGGGGGSSSSAG    290
           *:. :  ****:*:. ************** :***:     .**.  **       .. .
                                              BELL domain
           SSKDVPTLSAADRIEHQRRKVKLLSMLDEVDRRYNHYCEQMQMVVNSFDLVMGFGAAVPY    426
           SSKDVPTLSAADRIEHQRRKVKLLSMLDEVDRRYNHYCEQMQMVVNSFDLVMGFGAAVPY    426
           ANKEHPPLSASDRIEHQRRKVKLLTMLEEVDRRYNHYCEQMQMVVNSFDIVMGHGAALPY    424
           TANDSPPLSPADRIEHQRRKVKLLSMLEEVDRRYNHYCEQMQMVVNSFDQVMGYGAAVPY    350
           : :: * ** :*************:**:********************* ***.***:**

           TALAQKAMSRHFRCLKDAIGAQLKQSCELLGEKDA---GTSGLTKGETPRLKMLEQSLRQ    483
           TALAQKAMSRHFRCLKDAIGAQLKQSCELLGEKDA---GTSGLTKGETPRLKMLEQSLRQ    483
           TALAQKAMSRHFRCLKDAVAAQLKQSCELLGDKDAAGISSSGLTKGETPRLRLLEQSLRQ    484
           TTLAQKAMSRHFRCLKDAVAVQLKRSCELLGDKEAAGAASSGLTKGETPRLRLLEQSLRQ    410
           *:****************:..***:******:*:*   .:***********::*******
                                       Homeodomain
           QRAFHQMGMMEQEAWRPQRGLPERSVNILRAWLFEHFLHPYPSDADKHLLARQTGLSRNQ    543
           QRAFHQMGMMEQEAWRPQRGLPERSVNILRAWLFEHFLHPYPSDADKHLLARQTGLSRNQ    543
           NRAFHQMGMMEQEAWRPQRGLPERSVNILRAWLFEHFLHPYPSDADKHLLARQTGLSRNQ    544
           QRAFHHMGMMEQEAWRPQRGLPERSVNILRAWLFEHFLNPYPSDADKHLLARQTGLSRNQ    470
           :****:********************************:*********************

           VSNWFINARVRLWKPW--------------------------------------------    559
           VSNWFINARVRLWKPMVEDMYQQEAKDEDENSQSQNSGNNIIAQTPTPNS--LTNSSSTN    601
           VSNWFINARVRLWKPMVEEMYQQESKEREREEELEENEEDQETKNSNDDKSTKSNNNESN    604
           VSNWFINARVRLWKPMVEEMYQQEAKEREEAEEENENQQQQRRQQQTNNNDTKPNNNENN    530
           ***************

           ------------------------------------------------------------    559
           ITTATAAATTTTAPTTTTALAAAETGTAATPITVTSSKRSQINATDSDPSLVA-INSFSE    660
           FTA---VRTTSQTPTTTAPDASD----ADAAVATGHRLRSNINAYENDASSLLLPSSYSN    657
           FTV---ITAQT--PTTMTSTHHE-----------------------NDS-SF--------    553


           ------------------------------------------------------------    559
           NQATF--T----------TNIHDPDACRRGNFSGDDGTTTHDHMGSTMIRFGTT-AGDVS    707
           AAAPAAVSDDLNSRYGGSDAFSAVATCQQS----VGGFDDADMDGVNVIRFGTNPTGDVS    713
           ------LSSVAAASHGGSDAF-TVATCQQD----VSDFH-VDGDGVNVIRFGTKQTGDVS    601


           --------------------------    559
           LTLGLRHAGNLPENTHFFG-------    726
           LTLGLRHAGNMPDKDASFCVREFGGF    739
           LTLGLRHSGNIPDKNTSFSVRDFGDF    627
```

**Supplementary figure 6.** Alignment of BIP and other BLH proteins.

Identical or similar amino acid residues were indicated by asterisks and colons, respectively. Alignment was performed with CLUSTALW.

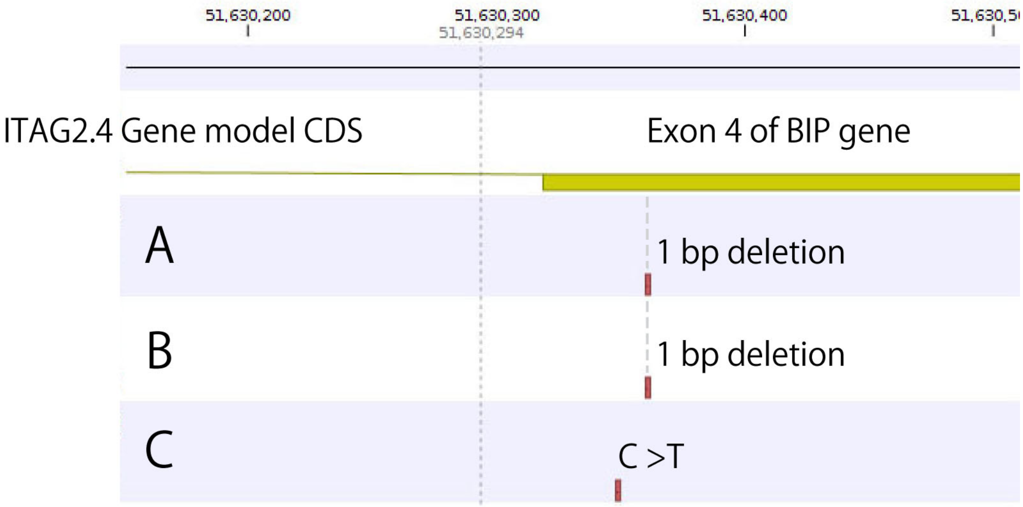


**Supplementary figure 7.** Mutations in *BIP* gene in different genomes.

(A) A SNP in the SiFT genome sequenced in this study. (B) A SNP in another SiFT genome sequenced by Tieman and coworkers in Tieman D, *et al*. (2017): A chemical genetic roadmap to improved tomato flavor. Science 355(6323):391-394. (C) A SNP in *bip3* mutant genome. Mapping and SNP calling were performed with CLC genomic workbench, then data was visualized.

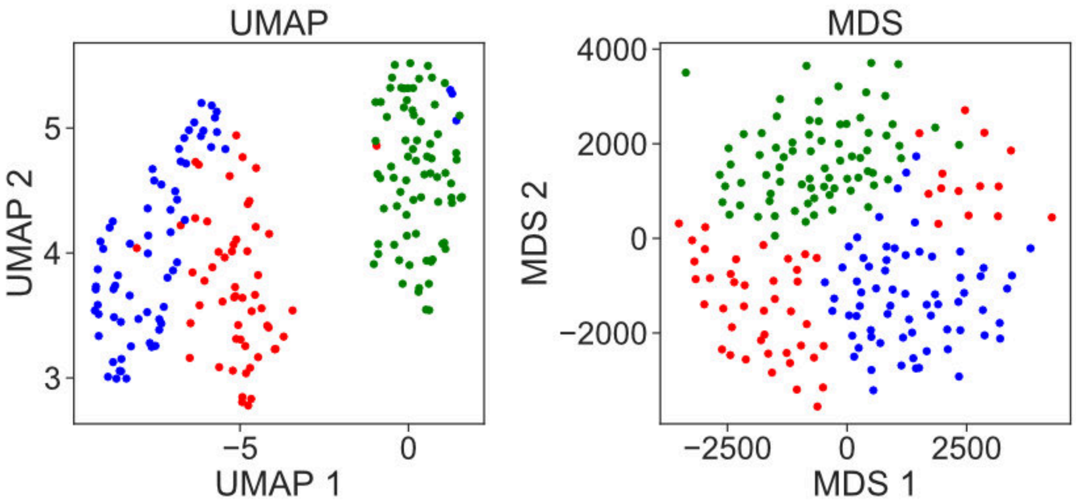


**Supplementary figure 8.** Machine learning-based leaflet shape analysis.

Results of UMAP (left), and MDS (right) with a pre-trained neural network with the ImageNet dataset, VGG19 (N < 55). The same data set shown in Figure 3F was used. Green: M82; Red: *bip3*; Blue: SiFT.

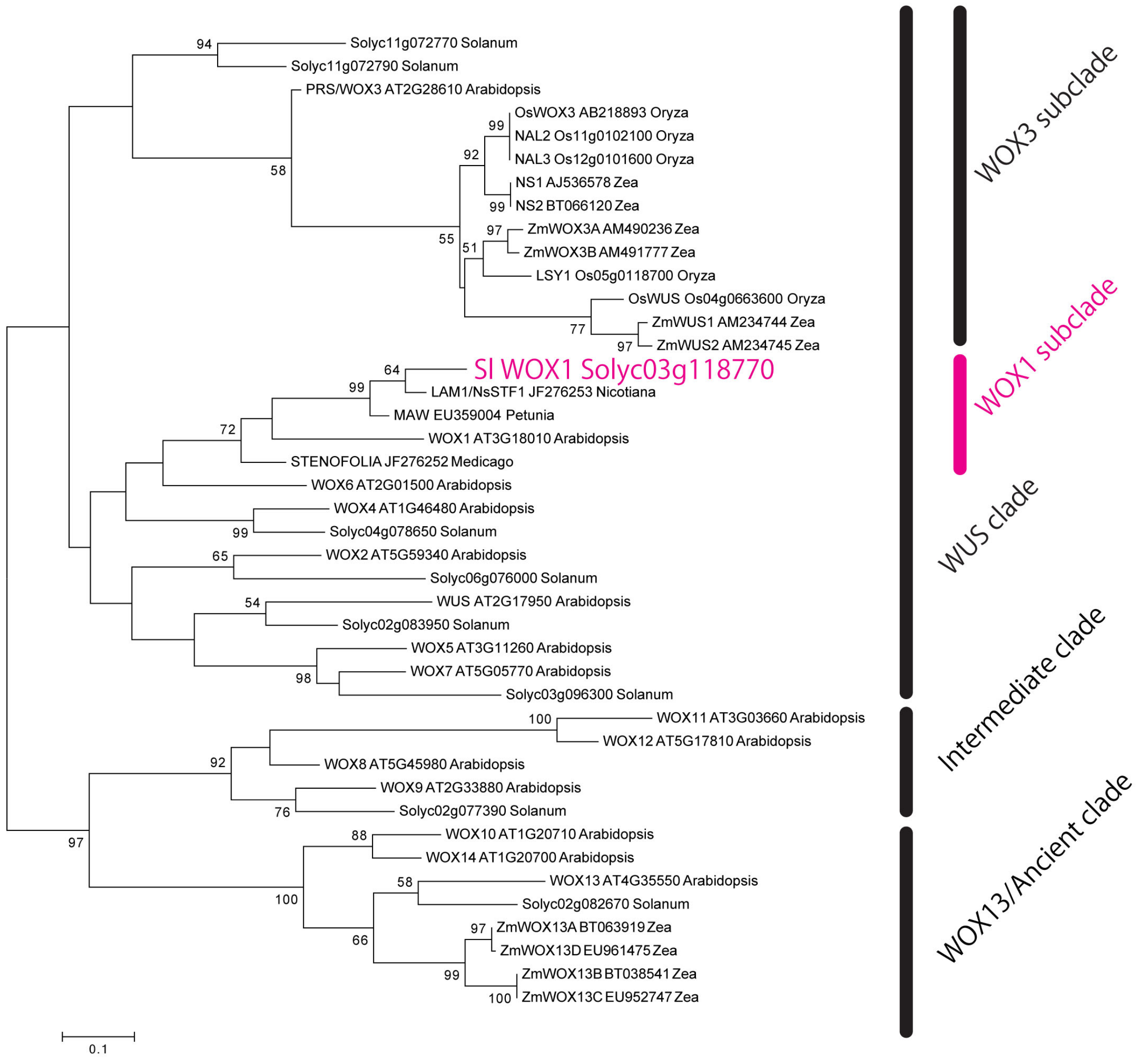


**Supplementary figure 9.** A phylogenetic tree of *WOX* orthologs.

A phylogenetic tree of *WOX* orthologs resulting from Neighbor-joining method by MEGA. Sl *WOX1* is located in a *WOX1* subclade in the phylogeny. The bootstrap values are indicated on branches (only those more than 50% are indicated on the tree).

```
SlWOX1_Tomato            MWMMGYNEGGDFNMSDS-CNGRKLRPLMPR---VPHAPTANPTN---CLRNFHGENFIAL
LAM1/NsSTF1_Tobacco      MWMVGYNDGGDFNMQDS-FNGRKLRPLMPR---VPHLPTANISTNPTCLRSIHGENFVAL
STENOFOLIA_Medicago      MWMVGYNEGGEFNMADYPFSGRKLRPLIPRPVPVPTTSPNNTSTITPSLNRIHGGNDLFS
WOX1_Arabidopsis         MWTMGYNEGG----ADSFNGGRKLRPLIPR---LTSCPTAAVNTNSDHRFNMAVVTMTAE
                         ** :***:**     *   .*******:**   :.  ..   ..       :   .
                                                              Homeodomain
                         NHHQ-----------LAMSEQNKRDFN-TQLVVSSRWNPTPEQLQTLEELYRRGTRTPSA
                         NHHQ-----------LAMSEQNKRDFNTQQLVVSSRWNPTPEQLQTLEELYRRGTRTPSA
                         QYHHNLQQQASVGDHSKRSELNNNNNPSAAVVVSSRWNPTPEQLRALEELYRRGTRTPSA
                         QNKR-------------ELMMLNSEPQHPPVMVSSRWNPTPDQLRVLEELYRQGTRTPSA
                         : ::                  : :     ::*********:**:.******:*******

                         EQIQHITAQLRRYGKIEGKNVFYWFQNHKARERQKRRQLESSANGNGN--GGGGDDQSQ
                         EQIQHITAQLRRYGKIEGKNVFYWFQNHKARERQKRRQLESAAGGGAANAAGGGDDQSR
                         EQIQQITAQLRKFGKIEGKNVFYWFQNHKARERQKRRQMESAAAEFDS-----------
                         DHIQQITAQLRRYGKIEGKNVFYWFQNHKARERQKRRQMETGHEETVLS----------
                         ::**:******::**************************:*:.

                         SNCNAENAERKESGANRTVFEIEQTKHWPSPTNCSTLAEKTAAKTKAGAASAAAGATAAG
                         SNCNPENTERKESGANRTGFEIEQTKNWPSPTNCSTLAEKTVATTKAAAAGGVAECRVA-
                         ------AIEKKDLGASRTVFEVEHTKNWLPSTNSSTSTLPLAEESVSIQRSAAAK-----
                         --TASLVSNHGFDKKDPPGYKVEQVKNWICSVGCDTQPEKPSRDYHLEEP----------
                                 ::     . . :::*:.*:*  .....* .

                         VAESCRVAAAERWIPFDEGEQRRSLLAER--NATWQMMHLSCSPPTIN-------NNTNC
                         ---------AERWIPFDEGEQRRSLLLADQRNATWQMMHLSCSPPTSTTPHHHLMNINSS
                         ---------ADGWLQFDEAELQQRRNFMER-NATWHMMQLTSSCPTAS------------
                         ---------ANIRVEHNARCGGDERRSFLGINTTWQMMQLPPSFYSSS--------HHHH
                                  *:  : .:              *:**:**:*. *  : .

                         ATICSNTITTATCTPIIRSCPSTPTTIDHQTKQLFKPKDHLNIFITPFRCDQKHQNIIGD
                         TAAISNTISGTTSPIICSSNTPSTPRTTMEPKQLFKTKDHLNIFIAPFRTDNRKHENMEN
                         -------------------MSTTTTVTTRLMDPKLIKTHELNLFISPHTYKERENAFIHL
                         QRNLILNSPTVSSNMSNSNNAVSASKDTVTVSPVFLRTREATNTETCHRNGDDNKDQEQH
                                             . :..      .  :  . . .   : .   : .:
                                   WOXdomain
                         EEEEGEGNGHEAQTLELFPLRSSNDNNDENNFSDKD---EIGAAANLNNNFNGSHYQFFE
                         --IVGDEGQEESQTLELFPLRSSNDNNDDNNFSEKDEVEISGADANSNSNFSGSHCQFFE
                         N-TSSTHQNESDQTLQLFPIRNGDHGCTDHHHHHHNIIKETQISASAINAPN----QFIE
                         EDCSNGELDHQEQTLELFPLRKEGFCSD------------------GEKDKNISGIHCFYE
                             .    .. ***:***:*. .                     .  .        * *

                         FLPLKN
                         FLPLKN
                         FLPLKN
                         FLPLKN
                         ******
```

**Supplementary figure 10.** Alignment of Sl WOX1 and other WOX1 proteins.

Identical or similar amino acid residues were indicated by asterisks and colons, respectively. Alignment was performed with CLUSTALW.

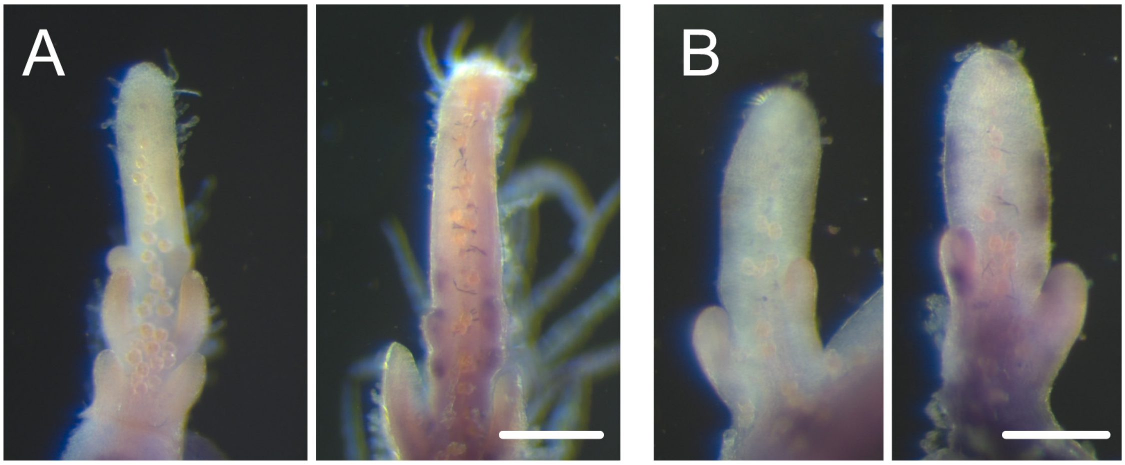


**Supplementary figure 11.** Expression pattern of Sl *WOX1* in SiFT.

*in situ* localization of Sl *WOX1* transcripts in SiFT. (A) Leaf primordia. (B) Leaflet primordia. Left: sense probe; right: antisense probe in each panel. Bars = 100 $\mu$m.

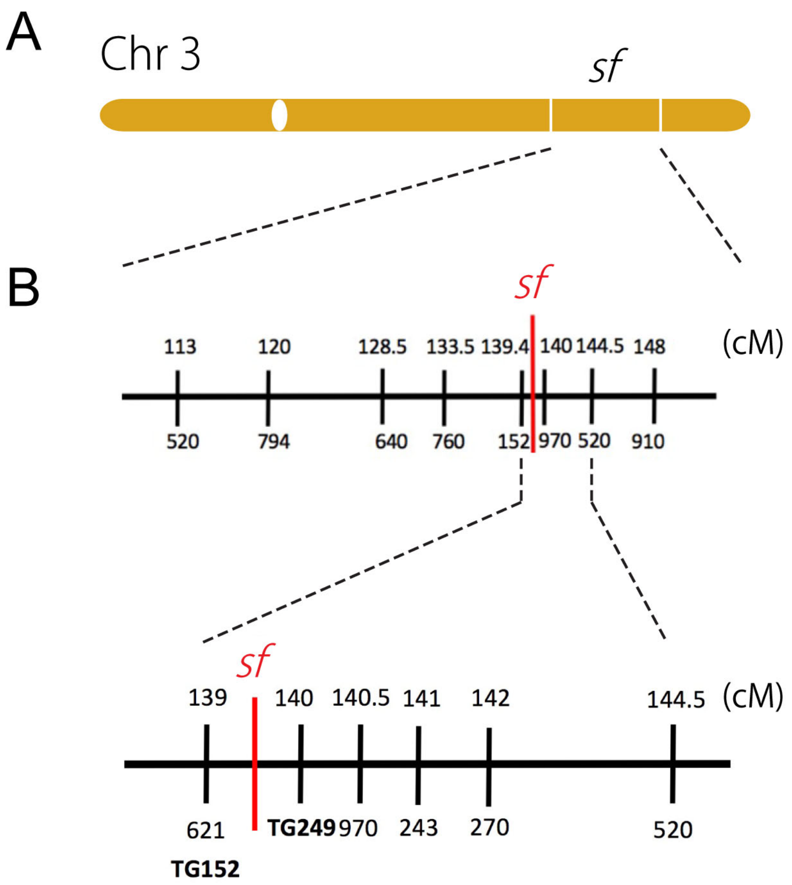


**Supplementary figure 12.** Map-based cloning of *sf*.

(A) Location of *sf* on chromosome 3. (B) Rough mapping of *sf* locus. Numbers below and above the map indicate makers and the physical distance between markers, respectively.

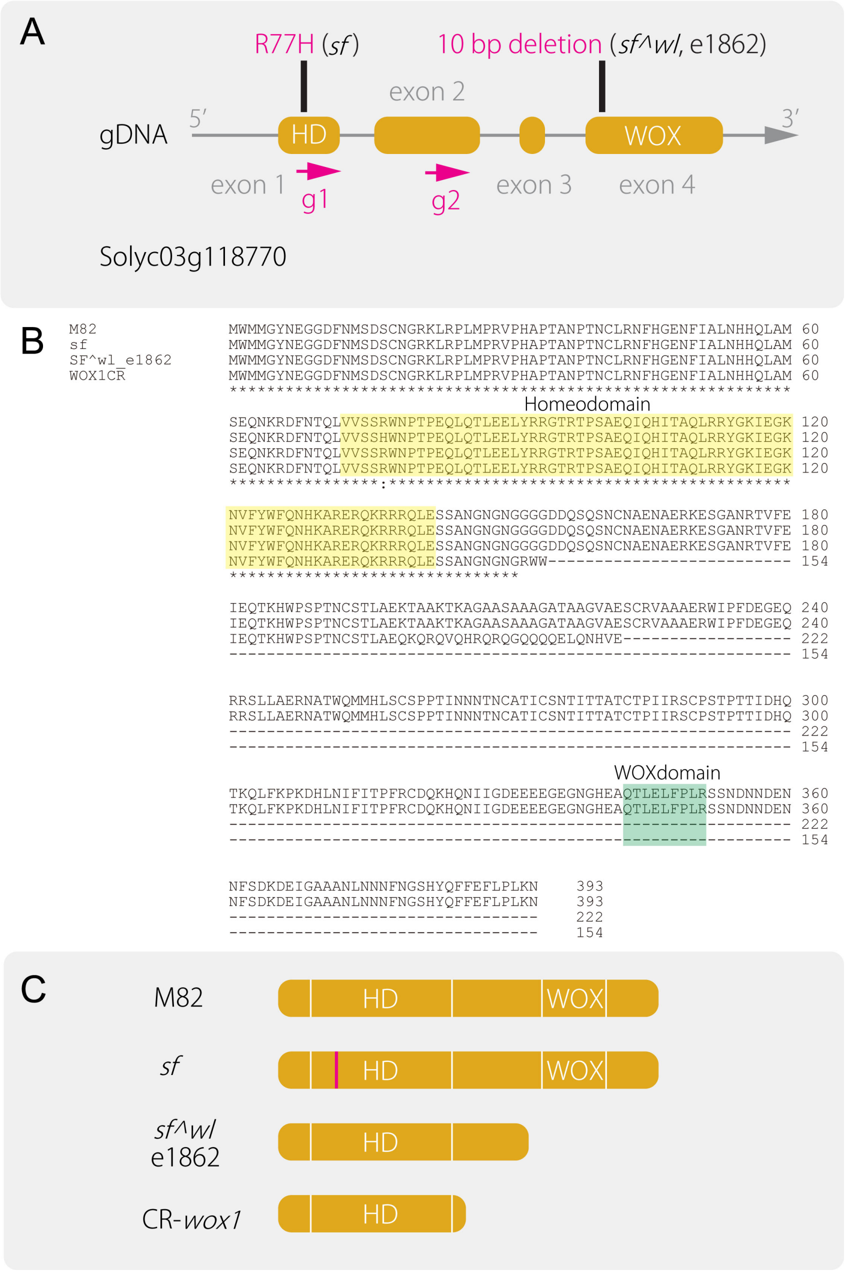


**Supplementary figure 13.** Comparison of *wox1* mutant alleles.

(A) Exon and intron structure of *SOLANIFOLIA*/*SLWOX1*(*sf*/*slwox1*). The tomato *SF*/*SLWOX1* gene contains four exons. (B) Alignment of Sl WOX1 and other WOX1 proteins. Identical or similar amino acid residues were indicated by asterisks and colons, respectively. Alignment was performed with CLUSTALW. (C) Comparison of proteins from different *wox1* mutant alleles.

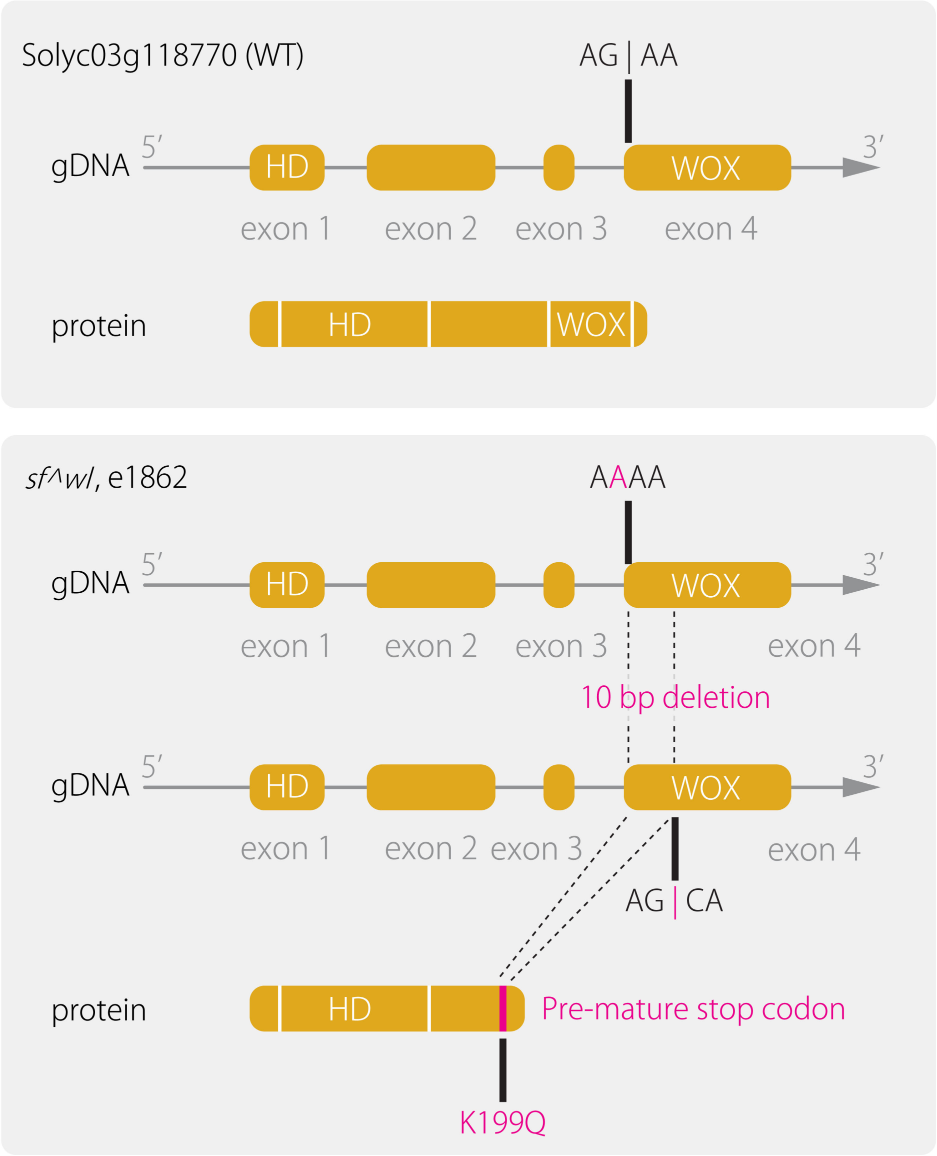


**Supplementary figure 14.** WOX1 proteins in *sf^wl* and e1862 lack the conserved WOX domain.

Exon and intron structure of M82 (Top; WT) and *sf^wl* and e1862 (bottom). M82 has a splice site (vertical line) at AG between intron 3 and exon 4. *sf^wl* and e1862 have a 1 bp substitution (G to A) at the splice site, which results in 10 bp shift to the next splice site. As a result, these mutants have a 10 bp deletion from position 595 to 604 resulting in a premature stop codon, which truncates the WOX1 protein such that it lacks the conserved WOX domain.

```
SFT     ATGTGGATGATGGGTTACAATGAGGGAGGAGATTTTAACATGTCGGATTCGTGTAATGGA     60
M82     ATGTGGATGATGGGTTACAATGAGGGAGGAGATTTTAACATGTCGGATTCGTGTAATGGA     60
        ************************************************************

        AGAAAGCTTCGTCCACTAATGCCGAGAGTGCCTCATGCGCCTACGGCGAATCCAACGAAT     120
        AGAAAGCTTCGTCCACTAATGCCGAGAGTGCCTCATGCGCCTACGGCGAATCCAACGAAT     120
        ************************************************************

        TGCTTAAGAAATTTTCATGGAGAAAACTTTATTGCACTTAATCATCATCAGCTTGCTATG     180
        TGCTTAAGAAATTTTCATGGAGAAAACTTTATTGCACTTAATCATCATCAGCTTGCTATG     180
        ************************************************************

        AGTGAGCAAAATAAGAGAGATTTCAATACGCAATTAGTAGTGAGCTCACGTTGGAATCCA     240
        AGTGAGCAAAATAAGAGAGATTTCAATACGCAATTAGTAGTGAGCTCACGTTGGAATCCA     240
        ************************************************************

        ACTCCAGAACAACTGCAAACGCTGGAAGAGTTGTATCGACGCGGCACGAGAACTCCGTCA     300
        ACTCCAGAACAACTGCAAACGCTGGAAGAGTTGTATCGACGCGGCACGAGAACTCCGTCA     300
        ************************************************************

        GCTGAACAGATTCAGCATATCACTGCACAACTTAGACGGTATGGCAAAATTGAAGGAAAA     360
        GCTGAACAGATTCAGCATATCACTGCACAACTTAGACGGTATGGCAAAATTGAAGGAAAA     360
        ************************************************************

        AATGTTTTCTACTGGTTTCAAAATCATAAAGCAAGGGAACGCCAAAAAAGACGCCGTCAA     420
        AATGTTTTCTACTGGTTTCAAAATCATAAAGCAAGGGAACGCCAAAAAAGACGCCGTCAA     420
        ************************************************************

        CTTGAATCTTCTGCTAATGGGAACGGTAATGGCGGTGGTGGTGATGATCAGTCTCAGAGT     480
        CTTGAATCTTCTGCTAATGGGAACGGTAATGGCGGTGGTGGTGATGATCAGTCTCAGAGT     480
        ************************************************************

        AATTGTAACGCTGAAAATGCTGAAAGAAAAGAATCAGGGGCAAATAGGACAGTTTTTGAA     540
        AATTGTAACGCTGAAAATGCTGAAAGAAAAGAATCAGGGGCAAATAGGACAGTTTTTGAA     540
        ************************************************************

        ATTGAACAGACCAAGCACTGGCCATCCCCAACAAACTGCAGTACTCTTGCAGAGAAAACT     600
        ATTGAACAGACCAAGCACTGGCCATCCCCAACAAACTGCAGTACTCTTGCAGAGAAAACT     600
        ************************************************************

        GCAGCAAAAACAAAGGCAGGTGCAGCATCGGCAGCGGCAGGGGCAACAGCAGCAGGAGTT     660
        GCAGCAAAAACAAAGGCAGGTGCAGCATCGGCAGCGGCAGGGGCAACAGCAGCAGGAGTT     660
        ************************************************************

        GCAGAATCATGTAGAGTAGCAGCAGCAGAAAGATGGATACCATTCGACGAAGGAGAACAA     720
        GCAGAATCATGTAGAGTAGCAGCAGCAGAAAGATGGATACCATTCGACGAAGGAGAACAA     720
        ************************************************************

        AGAAGGAGCTTATTAGCAGAAAGGAATGCCACGTGGCAGATGATGCATTTGTCTTGTTCA     780
        AGAAGGAGCTTATTAGCAGAAAGGAATGCCACGTGGCAGATGATGCATTTGTCTTGTTCA     780
        ************************************************************

        CCACCCACCATCAACAACAACACCAATTGTGCAACAATTTGTAGTAATACTATAACTACT     840
        CCACCCACCATCAACAACAACACCAATTGTGCAACAATTTGTAGTAATACTATAACTACT     840
        ************************************************************

        GCAACTTGTACTCCAATTATAAGGAGTTGTCCATCAACGCCAACAACAATCGACCATCAG     900
        GCAACTTGTACTCCAATTATAAGGAGTTGTCCATCAACGCCAACAACAATCGACCATCAG     900
        ************************************************************

        ACAAAACAACTTTTCAAGCCCAAAGATCATCTCAACATCTTTATAACACCTTTTAGATGT     960
        ACAAAACAACTTTTCAAGCCCAAAGATCATCTCAACATCTTTATAACACCTTTTAGATGT     960
        ************************************************************

        GATCAAAAACACCAAAACATCATCGGAGACGAAGAAGAAGAAGGCGAAGGCAATGGTCAT     1020
        GATCAAAAACACCAAAACATCATCGGAGACGAAGAAGAAGAAGGCGAAGGCAATGGTCAT     1020
        ************************************************************

        GAAGCACAGACTCTTGAATTATTTCCCCTCCGTAGCAGCAATGACAACAATGATGAAAAT     1080
        GAAGCACAGACTCTTGAATTATTTCCCCTCCGTAGCAGCAATGACAACAATGATGAAAAT     1080
        ************************************************************

        AATTTTTCAGACAAGGATGAAATAGGAGCTGCCGCGAACTTGAATAATAATTTTAATGGC     1140
        AATTTTTCAGACAAGGATGAAATAGGAGCTGCCGCGAACTTGAATAATAATTTTAATGGC     1140
        ************************************************************

        AGTCATTATCAGTTTTTTGAGTTCCTTCCACTCAAGAAC     1179
        AGTCATTATCAGTTTTTTGAGTTCCTTCCACTCAAGAAC     1179
        ***************************************
```

**Supplementary figure 15.** Alignment of Sl WOX1 between M82 and SiFT.

Identical or similar amino acid residues were indicated by asterisks and colons, respectively. Alignment was performed with CLUSTALW.

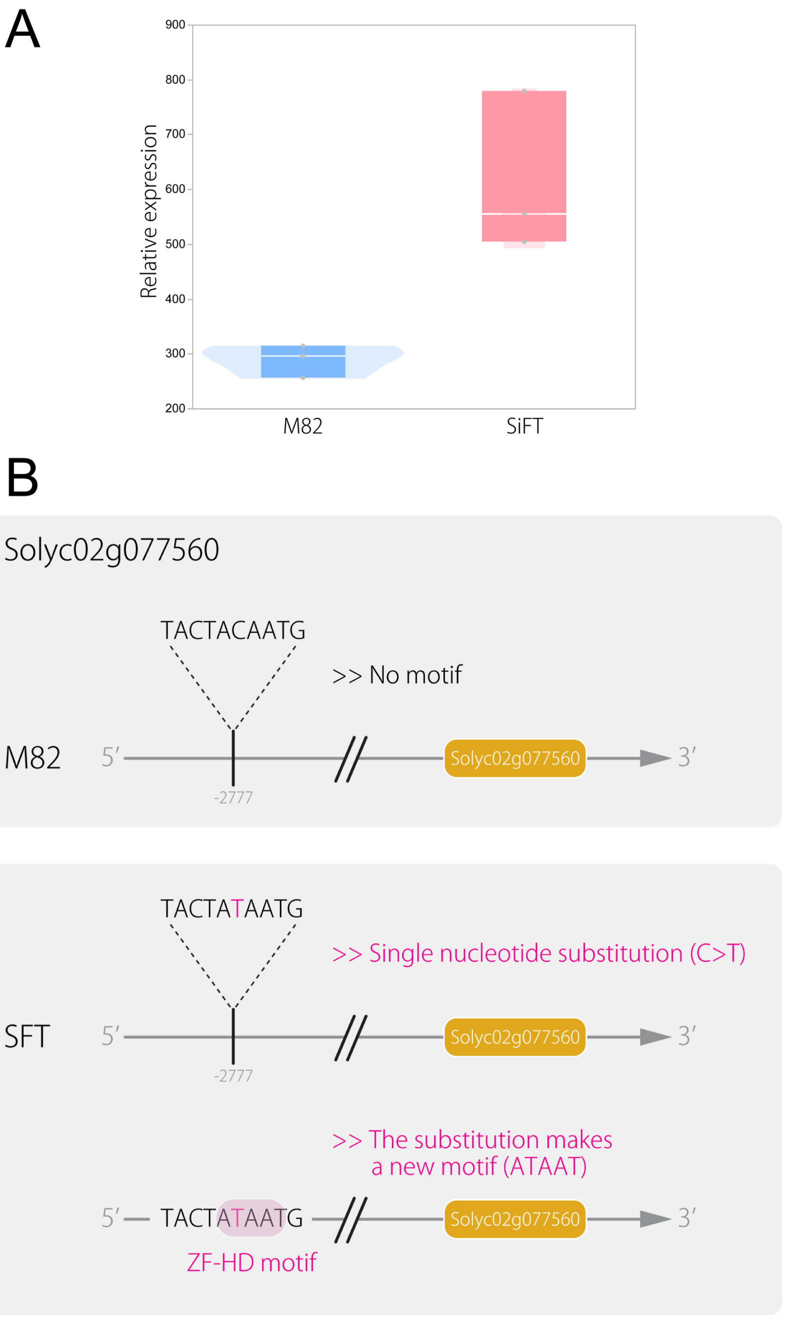


**Supplementary figure 16.** *ARF3* expression level and a difference in promoter sequence.

(A) Expression level of Sl *ARF3* in leaf primordia ($n$ = 3). (B) Comparison of promoter sequences between M82 and SiFT.

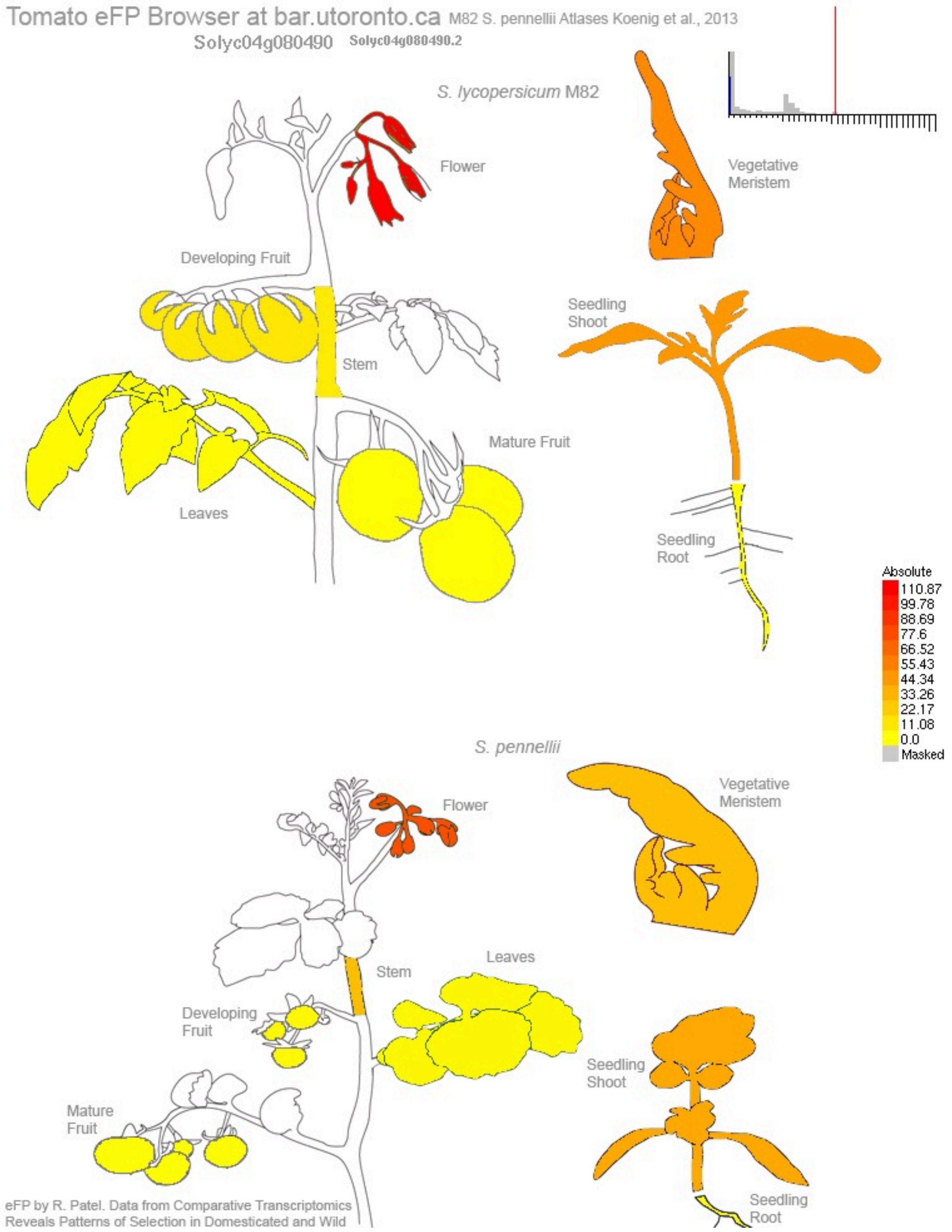


**Supplementary figure 17.** Expression pattern of Sl *HB33* in M82 and a wild species, *Solanum pennellii*.

Tomato eFP browser (http://bar.utoronto.ca/efp_tomato/cgi-bin/efpWeb.cgi
) shows expression of Sl *HB33* (Solyc04g080490) in organs of M82 (top) and organs in *S. pennellii* (bottom).

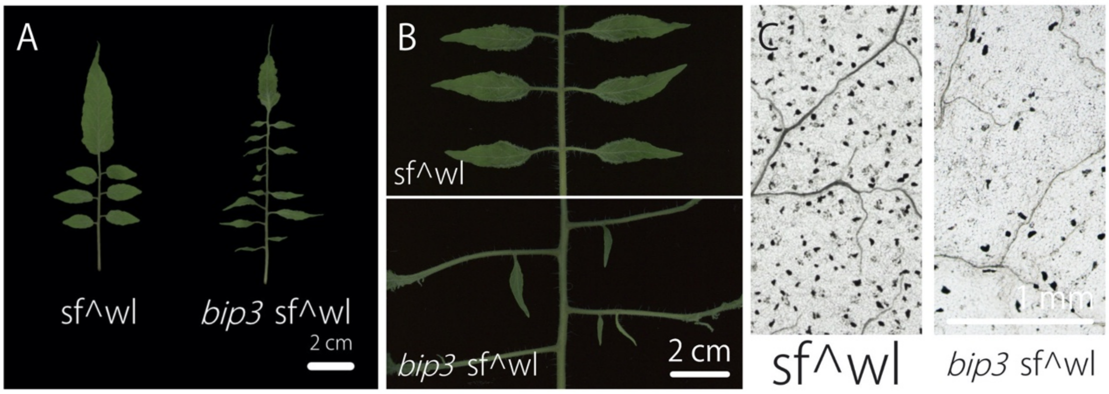


**Supplementary figure 18.** *bip sf/slwox1* leaf phenotypes.

(A) Mature leaf morphology of *bip3 sf^wl* double mutant. The 4th leaves were used. (B) Comparison of secondary leaflets on matured 6th leaf from 60 days old seedlings. (C) Cleared terminal leaflet images of *sf^wl* and *bip3 sf^wl* double mutant. Bars = 2 cm in (A) and (B), and 1mm in (C).

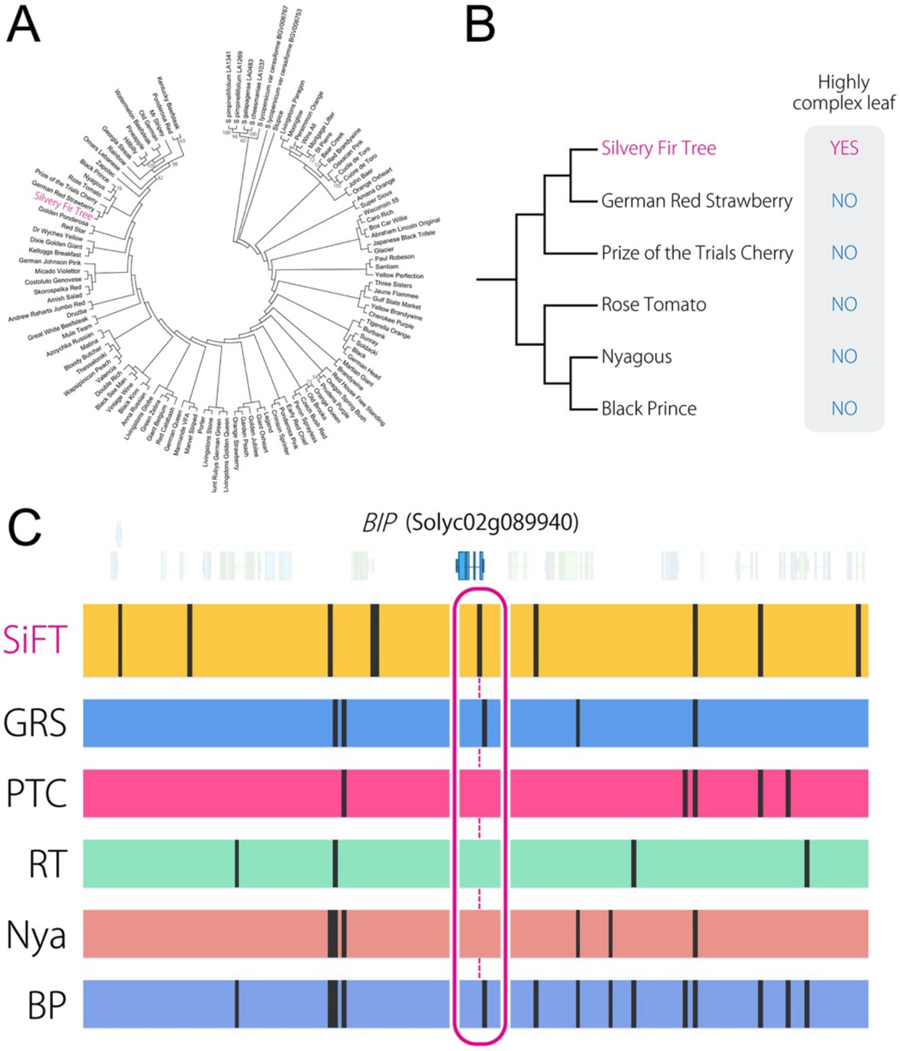


**Supplementary figure 19.** Phylogenetic tree with WGS data and comparison of SNPs data.

(A) ML Phylogenetic tree with whole genome sequencing data. The bootstrap values are indicated on branches (only those more than 50% are indicated on the tree). (B) Magnified view of the WGS phylogeny shown in (A) focusing on SiFT. (C) Comparison of SNPs data around *BIP* gene (Solyc02g089940) seen in SiFT. Each vertical black line indicates a SNP. German Red Strawberry (GRS), Prize of the Trials Cherry (PTC), Rose Tomato (RT), Nyagous (Nya), and Black Prince (BP). Note that a SNP seen in BP is not on the *BIP*.
